# Tissue-adapted NK cells shape pathogenic cDC1 niches in early arthritis

**DOI:** 10.64898/2026.05.01.716870

**Authors:** Oliver C. Knight, Chiara Giordano, Anne Sae Lim von Stuckrad, Cornelia Winning, Chrissy Bolton, Christopher B. Mahony, Mir-Farzin Mashreghi, Adam P. Croft, Tilmann Kallinich, Timo Rückert, Chiara Romagnani

## Abstract

Chronic forms of autoimmune arthritis, including juvenile idiopathic arthritis (JIA), are characterised by persistent synovial inflammation. While adaptive mechanisms are well-studied, the innate networks driving early joint disease remain poorly defined. Here, through unbiased profiling of innate lymphoid cells from the joints of children with JIA at disease onset, we identify activated natural killer (NK) cells as key orchestrators of adaptive immune niches. The selective upregulation of lymphotactin (XCL1) by activated NK cells is accompanied by the synovial enrichment of XCR1^+^ type 1 conventional dendritic cells (cDC1s), directly correlating with disease severity. In situ, NK cells spatially anchor with recruited cDC1s to assemble discrete, multicellular effector niches alongside T cells. Finally, we demonstrate this NK-cDC1 spatial architecture represents a conserved pathogenic module across adult arthropathies, including rheumatoid arthritis. Our findings establish the XCL1-XCR1 axis as a fundamental innate feature of chronic joint inflammation, defining a targetable mechanism that precedes and promotes downstream adaptive responses.

## Introduction

Juvenile idiopathic arthritis (JIA) is the most common childhood rheumatic disease, characterised by immune-mediated synovial inflammation that can result in chronic pain, restricted mobility, and irreversible joint damage^1^. Modern targeted therapies have dramatically improved clinical outcomes, yet sustained remission remains difficult to achieve for a substantial proportion of individuals. JIA shares core immunopathological features with adult arthropathies^2,3^, notably the synovial infiltration and clonal expansion of type 1 and type 17 CD4^+^ memory T cells and CD8^+^ effector memory T cells^4–8^. In adult conditions, upstream innate networks, such as macrophages and dendritic cells (DCs), are increasingly recognised as fundamental orchestrators of this adaptive pathology^9–11^. However, the contribution of innate lymphoid cells (ILCs) to the early stages of synovial inflammation remains poorly understood.

This compartment, comprising helper ILCs and natural killer (NK) cells, forms a heterogeneous lineage specialised for tissue homeostasis and early immunosurveillance^12^. Whereas human cytotoxic CD56^dim^ CD16^+^ (NK1) cells predominate in circulation, the CD56^bright^ CD16^lo/-^ (NK2) subset constitutes a minor fraction in blood^13^. Conversely, these cells are enriched in peripheral tissues^14^, where they exert immunoregulatory functions through cytokine secretion rather than direct cytotoxicity^15^. Flow cytometric analyses of JIA synovial fluid have shown that alongside helper ILCs^16^, NK cells constitute a substantial proportion (5-10%) of the mononuclear infiltrate^17^ and exhibit a primarily CD56^bright^-like phenotype^17,18^. Although the CD56^dim^/CD56^bright^ NK dichotomy is well established in blood^19–21^, unbiased transcriptional characterisations of tissue NK cell heterogeneity are only beginning to emerge^21–25^. It remains unknown whether these blood-derived paradigms adequately capture the true diversity of NK cells during chronic inflammation, and whether these joint-infiltrating cells reflect the simple recruitment of circulating CD56^bright^ cells or instead represent a distinct, locally adapted state with specialised functional programming.

Here, to resolve the identity of NK cells during autoimmune joint inflammation, we generate an unbiased innate lymphocyte atlas by integrating single-cell transcriptomics and surface protein expression from paired peripheral blood and synovial fluid samples of treatment-naïve JIA patients. We demonstrate that synovial NK cells undergo dynamic, proliferative adaptations to the local microenvironment, skewing their functional programme toward high expression of XCL1 and XCL2, the cognate ligands for XCR1. Alongside this localised chemokine production, we identify a marked enrichment of XCR1^+^ conventional type 1 dendritic cells (cDC1s) that correlates strongly with clinical pathology, mirroring the NK-cDC1 recruitment axis previously described in tumour microenvironments^26^. Using spatial transcriptomics, we further reveal that NK cells and cDC1s colocalise within discrete, T cell-centric effector niches in the inflamed synovium. Finally, we establish that this NK-cDC1 architecture represents a conserved spatial module across chronic adult arthropathies, including rheumatoid arthritis (RA), thereby defining a fundamental, upstream innate pathway that drives joint inflammation.

## Results

### A single-cell atlas of innate lymphoid cells in JIA synovial fluid

To resolve the NK and ILC landscape at the onset of clinical disease, we generated a multimodal dataset of paired peripheral blood and synovial fluid samples from children with newly diagnosed JIA. We isolated the broad innate lymphoid compartment (Lin^-^ CD3^-^ CD7^+^; Methods) from four patients and analysed it using cellular indexing of transcriptomes and epitopes by sequencing^27^ (CITE-seq) to simultaneously capture transcriptional states and cell surface protein phenotypes (**Fig. 1a and Extended Data Fig. 1a, b**).

**Fig. 1:**
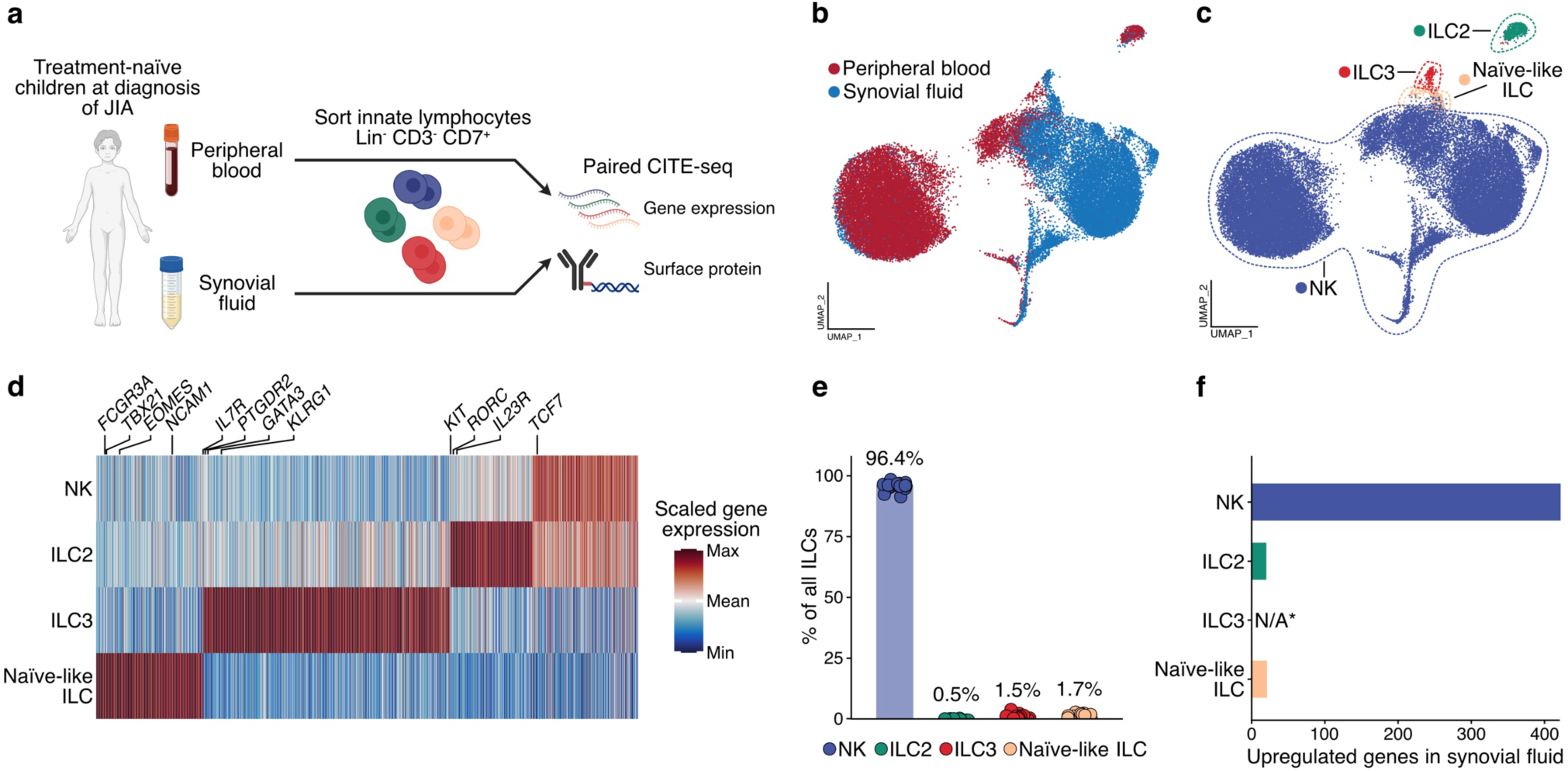
The transcriptional landscape of innate lymphoid cells in treatment-naïve JIA. **a**, Study design. Paired peripheral blood and synovial fluid samples were collected from juvenile idiopathic arthritis (JIA) patients at diagnosis (*n* = 4). Innate lymphoid cells (ILCs; Lin^-^ CD3^-^ CD7^+^) were isolated via fluorescence-activated cell sorting (FACS), pooled at a 1:1 ratio by origin, and analysed using CITE-seq. **b**, **c**, Integrated uniform manifold approximation and projection (UMAP) visualisations of annotated natural killer (NK) and helper ILC subsets (38,080 cells total; 19,750 from blood and 18,330 from synovial fluid). Cells are coloured by tissue origin (**b**) and assigned cell type (**c**). **d**, Heatmap detailing row-scaled differentially expressed genes across ILC subsets, with selected lineage markers highlighted. **e**, Frequencies of synovial fluid ILC subsets, expressed as a percentage of the total annotated ILC population. **f**, Number of differentially expressed genes enriched in synovial fluid compared to blood, categorised by cell type. Data in **e** and **f**, alongside all subsequent analyses, derive from our cohort integrated with a published CITE-seq dataset of paired blood and synovial fluid mononuclear cells from treatment-naïve patients with JIA^33^ (total *n* = 16). Bars indicate the mean ± s.d. Statistical significance was determined using a two-sided Wilcoxon rank-sum test with Bonferroni correction (**d**) and a paired Wald test via DESeq2 using blood as the reference (**f**). *N/A indicates that differential expression for ILC3s could not be calculated due to their absence in blood.

Unsupervised clustering of 38,080 recovered cells (**Fig. 1b)** identified 10 distinct transcriptional states across the patient cohort (**Extended Data Fig. 1c**), which we grouped and annotated using lineage-defining transcription factor and surface marker expression^28^ **(Fig. 1d and Extended Data Fig. 1d**). Within the helper ILC pool, we identified ILC2s (cluster 7, defined by *IL7R*, *GATA3, KLRG1* and CD294/CRTH2), ILC3s (cluster 8, defined by *RORC, IL23R*, CD117/c-KIT and NKp44), and ILC precursors^29^ or naïve-like ILCs^30^ (clusters 9 and 10, defined by *TCF7, ITGAX* and CD117/c-KIT, while lacking *RORC*) **(Fig. 1d and Extended Data Fig. 1d**). We did not detect populations resembling canonical helper^31^ (*IL7R*^+^ *TBX21*^+^ *RORC*^-^ *EOMES^-^*) or intraepithelial^32^ (*IL7R*^-^*TBX21*^+^ *RORC*^-^ *EOMES^-^ NCR2*^+^) ILC1s in either blood or synovial fluid (**Fig. 1c and Extended Data Fig. 1d**). By contrast, the NK cell lineage dominated the synovial innate lymphocyte landscape. This lineage encompassed six distinct clusters, unified by the expression of the core transcription factors *TBX21* and *EOMES*, alongside classical surface markers CD16 (*FCGR3A*) and CD56 (*NCAM1*) (**Fig. 1c, d and Extended Data Fig. 1c, d**).

To validate these findings, we integrated our dataset with an independent CITE-seq cohort^33^ of total mononuclear cells from paired blood and synovial fluid of nine treatment-naïve patients with JIA. This confirmed highly concordant frequency distributions and lineage-specific gene expression profiles within the inflamed joint (**Extended Data Fig. 1e, f**). To facilitate community access and independent exploration, this single-cell atlas is available for interactive analysis via the CELLxGENE platform (see Data Availability).

Differential expression analysis across this integrated cohort revealed that, in addition to dominating the innate lymphoid compartment in relative abundance (**Fig. 1e**), NK cells exhibited the most profound site-specific transcriptional divergence when compared to patient-matched blood. This population yielded the highest number of differentially expressed genes among all evaluated subsets (**Fig. 1f**). Given this pronounced tissue adaptation, we prioritised the NK cell compartment for downstream analyses, hypothesising that it serves as a central node of innate cellular remodelling during early joint inflammation.

### Synovial fluid NK cells undergo tissue-resident adaptation and activation

To understand how NK cells adapt to the inflamed joint in JIA, we compared their transcriptional profiles with matched blood counterparts (**Fig. 2a)**. Unlike circulating NK cells, cell cycle scoring placed a substantial fraction of synovial NK cells into S or G2/M phases (**Fig. 2a and Extended Data Fig. 2a**). We confirmed this proliferative signature by flow cytometry, finding a significantly higher frequency of Ki-67^+^ NK cells in synovial fluid relative to paired blood (**Fig. 2b**), identifying the inflammatory joint milieu as a potent mitogenic niche that drives local NK cell expansion.

**Fig. 2:**
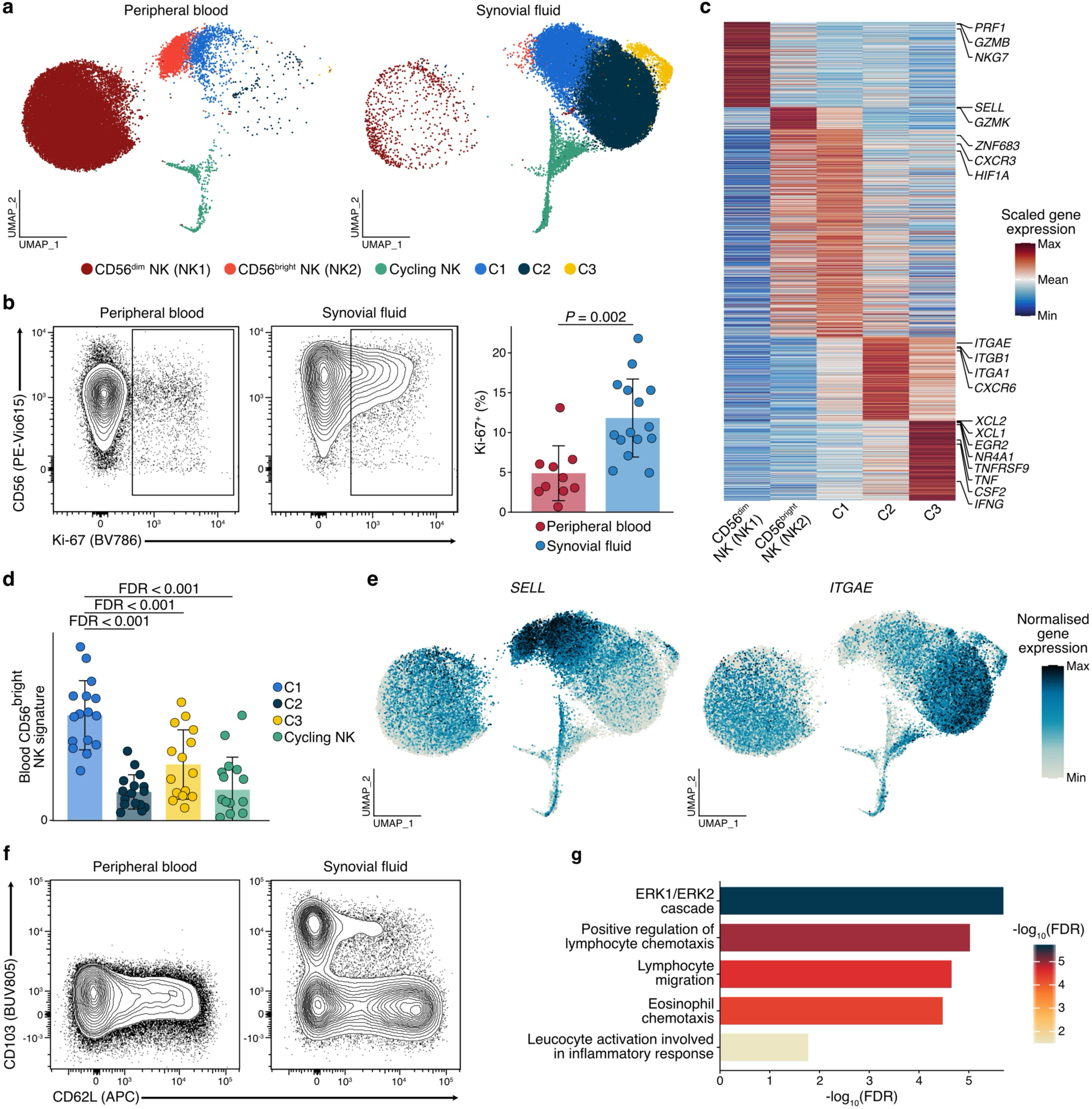
Phenotypic and transcriptional diversity of synovial fluid NK cells from JIA patients. **a**, Integrated UMAP of blood (30,451 cells) and synovial fluid (32,437 cells) NK cells from the complete annotated CITE-seq dataset. Cells are coloured by subpopulation. **b**, Representative flow cytometry plots (left) and quantification (right) of Ki-67 expression within the total NK cell population (gated as Lin^-^ CD3^-^ CD7^+^ EOMES^+^ NKp46^+^ events) from blood (*n* = 10) and synovial fluid (*n* = 15). Samples include 10 patient-matched pairs and 5 unpaired samples. **c**, Heatmap of row-scaled differentially expressed genes that define the annotated NK subpopulations, highlighting selected markers. **d**, Enrichment of the blood CD56^bright^ transcriptional signature across synovial NK subpopulations. **e, f,** Expression of *SELL*/CD62L and *ITGAE*/CD103 in paired blood and synovial fluid NK cells. Normalised gene expression is shown via CITE-seq feature plots (**e**), alongside corresponding surface protein expression demonstrated by representative flow cytometry plots (**f**). **g**, Top Gene Ontology Biological Process terms enriched among genes upregulated in the C3 subpopulation. Bars represent the mean ± s.d. Statistical significance was evaluated using a linear mixed-effects model with patient assigned as a random effect (**b**), a two-sided Wilcoxon rank-sum test with Bonferroni correction (**c**), paired two-tailed *t*-tests with Benjamini-Hochberg correction (**d**), and a hypergeometric test with Benjamini-Hochberg adjustment (**g**).

Peripheral blood NK cells segregated into the cytotoxic CD56^dim^ CD16^+^ (NK1) and immunoregulatory CD56^bright^ CD16^lo/-^ (NK2) subsets, distinguished by their canonical expression of *PRF1* and *GZMB* versus *SELL* and *GZMK*, respectively (**Fig. 2a, c and Extended Data Fig. 2b**). By contrast, only a minority of synovial NK cells mapped to these canonical circulating subsets, instead resolving into transcriptionally distinct states indicative of progressive tissue adaptation (**Fig. 2a, c**). The first synovial cluster (C1) largely mirrored the profile of recently infiltrated CD56^bright^ cells: alongside an enrichment of the blood CD56^bright^ NK signature (**Fig. 2d**), C1 maintained the expression of homing receptors *SELL* and *CXCR3* (**Fig. 2e**). However, when compared to their circulating CD56^bright^ counterparts, C1 cells were defined by the innate residency-associated transcription factor *ZNF683* (Hobit^34^) and hypoxia-inducible factor *HIF1A* (**Fig. 2c**). This suggests an early commitment to tissue retention. Conversely, the second cluster (C2) exhibited a definitive tissue-resident phenotype, characterised by expression of *ITGAE* (CD103), *ITGA1* (CD49a) and *CXCR6* (**Fig. 2c, e**). Using flow cytometry, we validated the coexistence of these distinct infiltrating (C1; CD62L⁺ CD103⁻) and tissue-resident (C2; CD62L⁻ CD103⁺) populations within the joint (**Fig. 2f and Extended Data Fig. 2c**). This phenotypic bifurcation was preserved within the Ki-67^+^ pool, indicating that in situ expansion occurs across both stages of adaptation (**Extended Data Fig. 2c**).

Finally, we identified a third synovial cluster (C3) representing a highly activated state. These cells were defined by the enriched expression of immediate-early genes^35^ (*EGR2*, *NR4A1*) and the effector cytokines *TNF*, *CSF2* (GM-CSF) and *IFNG* (IFN-γ) (**Fig. 2c**). As these cells concurrently expressed *ITGAE* and *CXCR6*, they map as an acutely activated state of the broader tissue-resident pool (**Extended Data Fig. 2d**). Gene set enrichment analysis (GSEA) revealed that alongside ERK1/ERK2 cascade signalling, the transcriptional programme of C3 was strongly enriched for pathways inducing lymphocyte migration and chemotaxis (**Fig. 2g**). This was driven by the high expression of chemokines *XCL1*, *XCL2*, *CCL3, CCL4* and *CCL5* (**Fig. 2c and Extended Data Fig. 2d, e**). Together, these findings indicate that the synovial microenvironment orchestrates NK cell expansion and tissue-resident reprogramming, pointing to a possible role in amplifying local immune cell recruitment through a distinct chemotactic signature.

### Synovial NK cells are the primary source of the cDC1 chemoattractant XCL1

The transcriptional signature of the highly activated synovial NK cell state (C3) was primarily defined by *XCL1* and *XCL2* (**Fig. 2c**). These paralogous lymphotactins serve as the principal chemotactic ligands for XCR1^+^ type 1 conventional dendritic cells (cDC1s)^36^, a myeloid subset specialised for antigen cross-presentation to CD8^+^ T cells^37,38^. Across the synovial immune landscape, *XCL1* and *XCL2* transcripts were predominantly expressed by the NK cells (**Fig. 3a**). Although memory CD8⁺ T cells can also contribute to the synovial *XCL1* pool^39^, NK cells were the primary cellular source of these chemokines within the inflamed joint (**Extended Data Fig. 3a**). This targeted production sharply contrasted with the expression of the canonical chemokine *CCL5*, which was shared with the CD8^+^ T cell compartment (**Fig. 3a**).

**Fig. 3:**
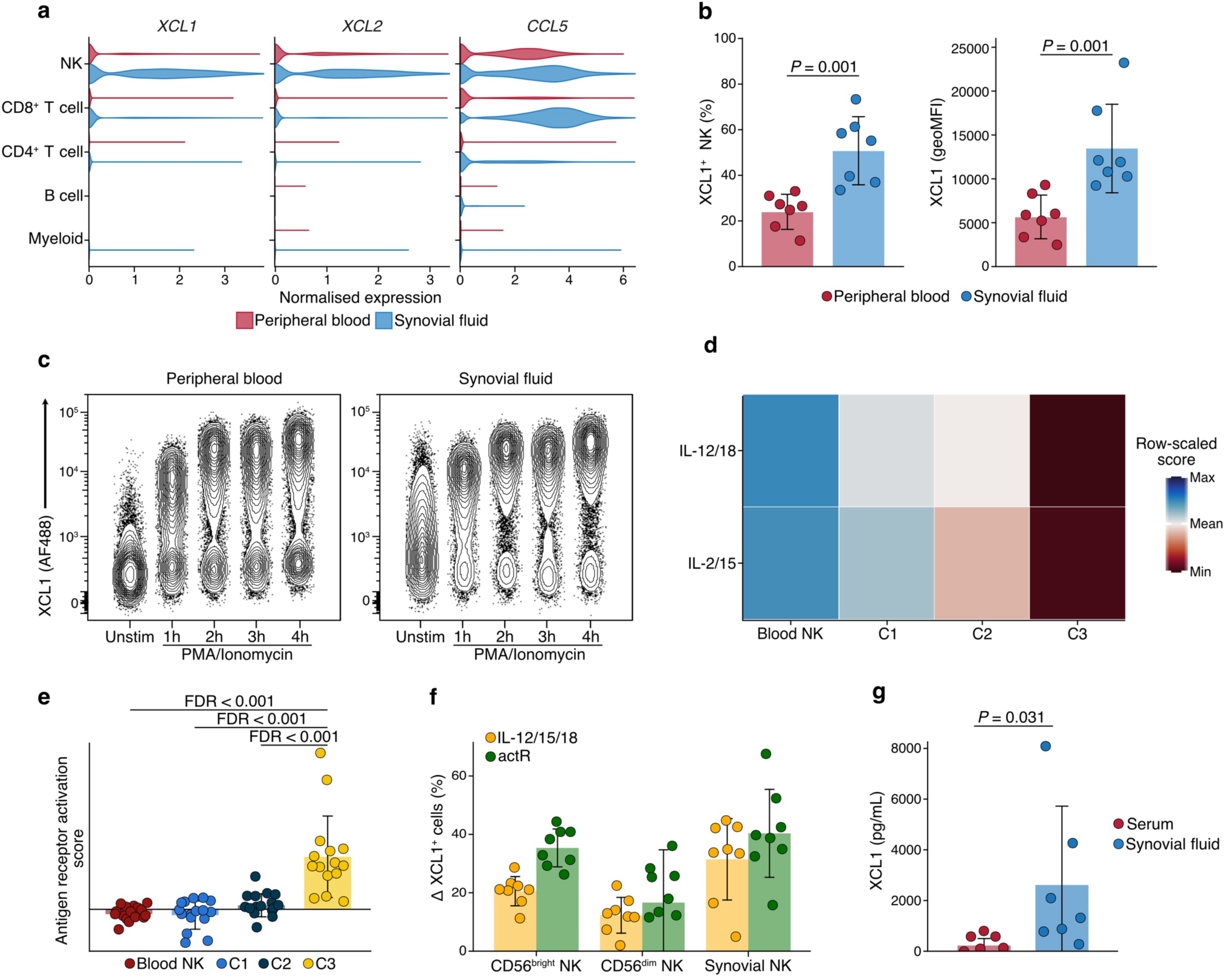
Synovial NK cells adopt a poised effector state characterised by robust XCL1 production. **a**, Violin plots detailing normalised *XCL1*, *XCL2,* and *CCL5* expression across annotated immune cell types in paired blood and synovial fluid (CITE-seq, *n* = 11). **b**, Frequencies (left) and geometric mean fluorescence intensities (geoMFI; right) of XCL1^+^ NK cells, calculated as a percentage of total NK cells (gated as live, Lin^-^ CD3^-^ CD7^+^ events) in blood (*n* = 8) and synovial fluid (*n* = 7; 7 paired, 1 unpaired) following a 6-hour unstimulated culture containing brefeldin A and monensin. **c**, Representative flow cytometry of intracellular XCL1 in total NK cells stimulated for up to 4 hours with PMA and ionomycin, with brefeldin A and monensin. **d**, **e**, Enrichment scores for transcriptional modules associated with cytokine-stimulated NK cells^40^ (**d**) and antigen-receptor T cell activation^41^ (**e**) within NK subpopulations. **f**, Absolute change in XCL1^+^ NK cell frequency relative to unstimulated controls in paired blood and synovial fluid (*n* = 8) following ex vivo stimulation with cytokines (IL-12, IL-15, and IL-18) or equimolar bead-conjugated activating receptor antibodies (actR; targeting CD2, CD226, CD244, NKG2D, NKp30, and NKp46). **g**, Quantification of XCL1 protein in paired serum and synovial fluid from patients with JIA, measured by ELISA (*n* = 7). Bars indicate the mean ± s.d. Statistical significance was determined using paired two-tailed *t*-tests (restricted to the 7 matched pairs for **b**), paired two-tailed *t*-tests with Benjamini-Hochberg correction (**d**), and a Wilcoxon signed-rank test following four-parameter logistic regression (**g**).

Given the high co-expression and functional redundancy^42^ of *XCL1* and *XCL2* (**Extended Data Fig. 3b**), we focused our protein-level validation on XCL1. Intracellular flow cytometry revealed that synovial fluid NK cells adopt a poised effector state. Compared with their circulating counterparts, they demonstrated robust and spontaneous production of XCL1 following six hours of ex vivo culture in the absence of exogenous stimulation (**Fig. 3b**). We did not observe this baseline activation in patient-matched CD8^+^ T cells (**Extended Data Fig. 3c**). Furthermore, this poised state was selectively skewed towards XCL1, as we observed no spontaneous production of the canonical, pro-inflammatory NK cell cytokines TNF and IFN-γ (**Extended Data Fig. 3d, e**). Consistent with this functional bias, stimulation with PMA and ionomycin induced a rapid upregulation of XCL1 that was detectable as early as one-hour post-stimulation (**Fig. 3c and Extended Data Fig. 3f**). Synovial NK cells also exhibited an increased capacity to produce TNF, but not IFN-γ, upon stimulation compared with blood NK cells (**Extended Data Fig. 3d, e**). This divergence further highlights the extensive functional reprogramming that occurs within the joint.

To gain insights into the possible stimuli driving XCL1 production, we considered previous studies indicating that cytokines can induce its expression in NK cells^43^. Guided by this, we applied module scoring using established activation signatures^40,41^. The activated synovial transcriptional state (C3) was significantly enriched for cytokine stimulation pathways, namely IL-2/IL-15 and IL-12/IL-18, as well as downstream antigen receptor activation (**Fig. 3d, e**). We functionally validated these transcriptomic predictions: targeted cytokine stimulation (IL-12, IL-15, and IL-18) and the specific cross-linking of multiple NK cell-activating receptors directly induced de novo XCL1 protein synthesis in both blood and synovial NK cells (**Fig. 3f**). Finally, we detected soluble XCL1 directly within patient synovial fluid aspirates, confirming its physiological secretion within the inflamed joint microenvironment (**Fig. 3g**).

Together, these findings establish the inflamed joint as a potent activating niche that drives NK cell-derived XCL1 production. This provides the necessary chemotactic gradient to recruit XCR1^+^ cDC1s to inflamed tissues, defining an innate immune axis previously described almost exclusively in the context of solid tumours^26^.

### The synovial microenvironment drives pathogenic cDC1 enrichment and functional adaptation

Prompted by the robust production of XCL1 by synovial NK cells, we investigated DC composition within the inflamed joint. Analysis of mononuclear cell CITE-seq data revealed a profoundly skewed synovial DC compartment compared with paired blood (**Fig. 4a and Extended Data Fig. 4a**). Although cDC2s remained the most abundant subset, the synovial fluid microenvironment was defined by a decreased frequency of plasmacytoid DCs (pDCs) and the emergence of *CCR7*^+^ mature regulatory DCs (mregDCs), a state primarily described in tumours^44,45^. We also observed a striking proportional expansion of synovial cDC1s, which contrasts with their typical scarcity in circulation (**Fig. 4a and Extended Data Fig. 4a**).

**Fig. 4:**
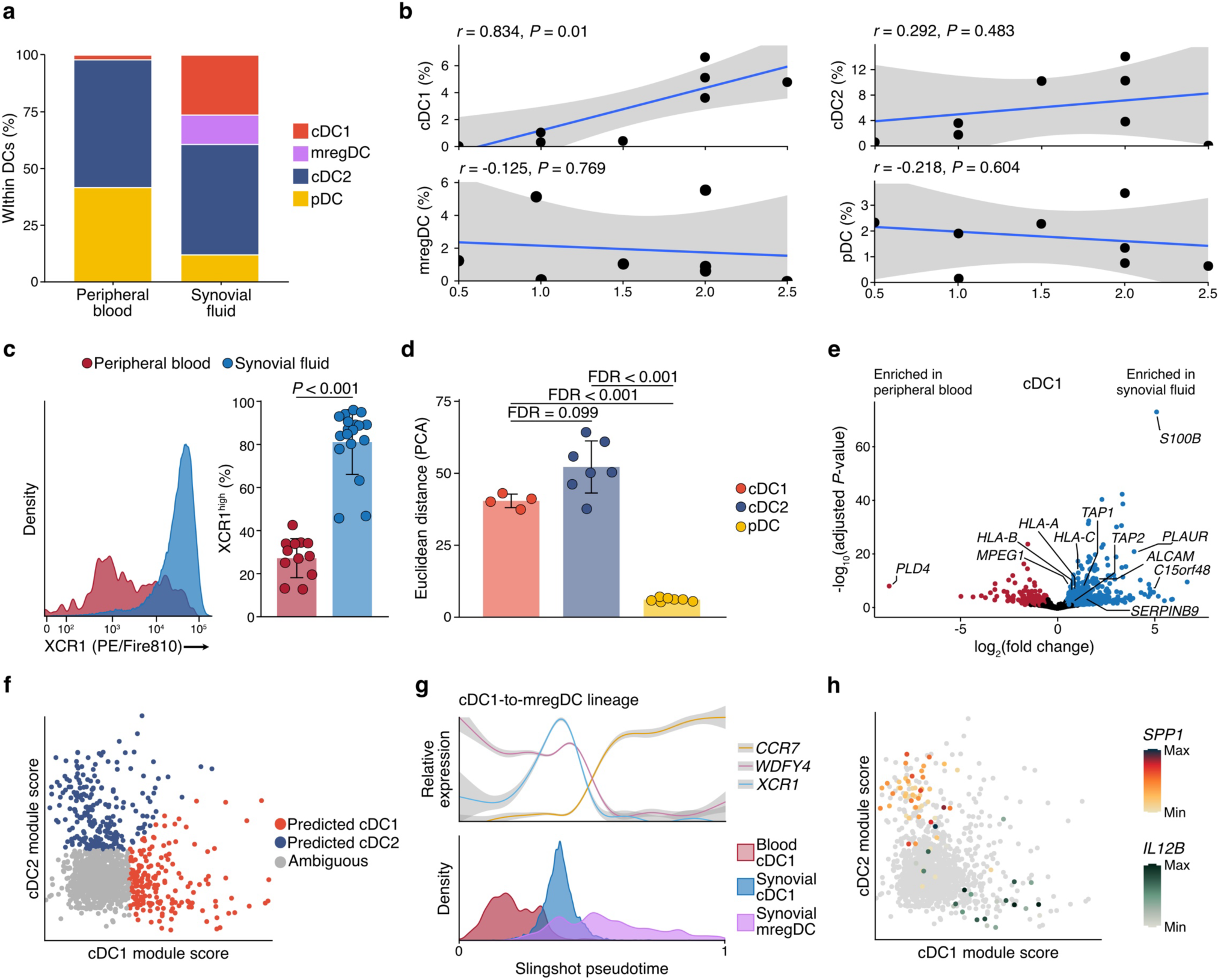
The transcriptional landscape of synovial fluid DC lineages in treatment-naïve JIA. **a**, Mean proportions of dendritic cell (DC) subsets in paired blood and synovial fluid, calculated as a percentage of total annotated mononuclear cells (CITE-seq, *n* = 11). **b**, Pearson correlation between the frequencies of synovial fluid DC subsets (expressed as a percentage of total annotated DCs) and the Krenn synovial lining hyperplasia score (CITE-seq, *n* = 8). The solid line indicates linear regression, and the shaded region denotes the 95% confidence interval (CI). **c**, Representative XCR1 distribution (left) and the frequency of XCR1^+^ cDC1s (right) in blood (*n* = 12) and synovial fluid (*n* = 18; 12 paired, 6 unpaired), as determined by flow cytometry. Frequencies are calculated as a percentage of total cDC1s (gated as live, Lin^-^CD11c^+^ CD141^+^ events). **d**, Mean Euclidean distance in principal component space between blood and synovial fluid DC subsets (CITE-seq; *n* = 4 for cDC1, *n* = 7 for cDC2, *n* = 7 for pDC, comprising 4 matched and 3 unpaired patients). **e**, Differentially expressed genes between cDC1s derived from blood and synovial fluid. **f, h,** Transcriptional module scoring and lineage characterisation of synovial fluid mregDCs, displaying predicted lineage origin (**f**) and the mutually exclusive expression of *IL12B* and *SPP1* (**h**; double-negative cells in grey). **g**, Slingshot pseudotime trajectory illustrating the transition from the cDC1 pool to mregDCs. The top panel displays the relative normalised expression of lineage-specific and maturation genes along the pseudotime axis; lines represent generalised additive model fits, with shaded areas indicating 95% CIs. The bottom panel depicts the relative density of cell populations across the inferred trajectory. Bars indicate the mean ± s.d. Statistical significance was assessed using a paired two-tailed *t*-test (restricted to the 12 matched pairs for **c**) with Benjamini-Hochberg correction (**d**), and a paired Wald test via DESeq2 using blood as the reference (**e**).

We evaluated these findings against available histopathological data^33^. Synovial fluid cDC1 abundance was the sole DC metric to correlate positively with the Krenn hyperplasia score, a measure of synovial tissue enlargement (**Fig. 4b**). This correlation strongly implicates the cDC1 subset in driving joint pathology. Consistent with targeted recruitment via the XCL1-XCR1 axis, transcript and flow cytometric analyses confirmed that synovial cDC1s expressed significantly higher levels of XCR1 compared to those in blood (**Fig. 4c and Extended Data Fig. 4b**). While previous studies in the tumour microenvironment have proposed that NK cell-derived CCL5 acts synergistically with XCL1 to recruit CCR5-expressing cDC1s^26^, the cognate receptor *CCR5* was notably absent on synovial cDC1s (**Extended Data Fig. 4b**), implicating synovial NK-derived XCL1 in their direct recruitment.

To map the functional specialisations of these subsets, we isolated pDCs (Lin^-^ CD303^+^), cDC1s (Lin^-^ CD303^-^ CD11c^+^ CD1c^-^ CD141^+^) and cDC2s (Lin^-^ CD303^-^ CD11c^+^ CD1c^+^) from both blood and synovial fluid and performing comparative CITE-seq profiling (**Extended Data Fig. 4c, d**; Methods). Global transcriptomic comparisons revealed that while synovial pDCs exhibited limited alterations compared with their circulating counterparts (**Extended Data Fig. 4e**), conventional DCs underwent extensive transcriptional adaptations to the inflamed joint (**Fig. 4d**). Both cDC subsets shared an expression programme characterised by the upregulation of canonical antigen presentation machinery, the inflammatory metabolic switch *C15orf48* (ref ^46^), and a common survival signature (**Fig. 4e and Extended Data Fig. 4f**). This included the granzyme B inhibitor *SERPINB9*, an upregulation likely required to maintain cellular integrity during close-contact cytotoxic interactions within the inflamed joint.

Beyond these shared pathways, the two canonical cDC subsets underwent distinct transcriptional reprogramming. Compared with blood, synovial cDC1s upregulated transcripts associated with trans-endothelial migration (*PLAUR*, *ALCAM*) alongside a pronounced signature for antigen cross-presentation, including a broad array of MHC class I molecules (*HLA-A, -B*, *-C*) and canonical processing machinery (*TAP1*, *TAP2*) (**Fig. 4e**). Within the synovial DC population, this cross-presentation signature was highly specific to cDC1s, marked by their exclusive expression of the essential cross-presentation genes *WDFY4* and *MPEG1* (ref. ^47,48^) (**Extended Data Fig. 4f**). By contrast, the adaptation of synovial cDC2s was driven by the acquisition of macrophage-associated tissue-remodelling genes^10,11^ (*MREG*, *CLEC5A*, *GPNMB*, *TREM2*), stromal activators (*FN1*, *PDGFB*) and an altered chemokine profile (*CCL22*, *CXCL16*) (**Extended Data Fig. 4g**).

Finally, we examined the synovial *CCR7*^+^ mregDC population. These cells were characterised by the expression of canonical markers^44^ (*LAMP3*, *FSCN1*, *BIRC3*), T cell-recruiting chemokines (*CCL19*, *CCL22*), and immunoregulatory checkpoint inhibitors (*CD274*, *PDCD1LG2*, encoding PD-L1 and PD-L2, respectively) (**Extended Data Fig. 4f**). Despite this uniform signature, module scoring utilising cDC1 and cDC2 gene sets revealed that synovial mregDCs derive from both lineages, maintaining discernible ancestral transcriptional footprints, alongside a substantial proportion of cells mapping to an ambiguous identity (**Fig. 4f**). Independent pseudotime analysis across the mregDC pool and both canonical cDC subsets confirmed that the acquisition of the mature mregDC state was marked by the induction of *CCR7* and coordinated loss of canonical cDC identifiers (**Fig. 4g and Extended Data Fig. 4h**). During this transition, cDC2s lost the lineage-defining marker *CD1C* and upregulated *MIR155HG*, a microRNA essential for driving mregDC differentiation^11^ (**Extended Data Fig. 4h**). In parallel, cDC1 maturation involved the simultaneous downregulation of both *XCR1* and *WDFY4* (**Fig. 4g**). Accordingly, mregDCs of distinct derivation exhibited divergent functional profiles: cDC2-derived mregDCs preferentially expressed the inflammatory factor *SPP1* (osteopontin), whereas cDC1-derived mregDCs specifically expressed *IL12B* (IL-12p40) (**Fig. 4h**). Together, these findings suggest a stepwise model wherein cDC1s are recruited to the inflamed joint via the XCL1-XCR1 axis and transcriptionally equipped for cross-presentation, before a subset subsequently transitions into a distinct mregDC state.

### NK cells spatially organise with cDC1s into discrete T cell niches in the synovium

To investigate the microanatomical distribution of these subsets in situ, we analysed spatial transcriptomic data from JIA patient synovial tissue^33^ to map the overarching cellular architecture across multiple patients (**Fig. 5a**). We first verified that global DC lineage signatures, alongside the expression of the key chemotactic receptor *XCR1*, were highly conserved between our synovial fluid samples and a published synovial tissue dataset^33^ (**Extended Data Fig. 5a, b**). Furthermore, the tight co-expression of the cognate NK cell-derived ligands, *XCL1* and *XCL2*, was maintained within the solid tissue (**Extended Data Fig. 5c**). Accordingly, the frequencies of both canonical cDC1s and mregDCs positively correlated with NK cell abundance within the tissue (**Fig. 5b**), an association we did not observe for canonical cDC2s (**Extended Data Fig. 5d**).

**Fig. 5:**
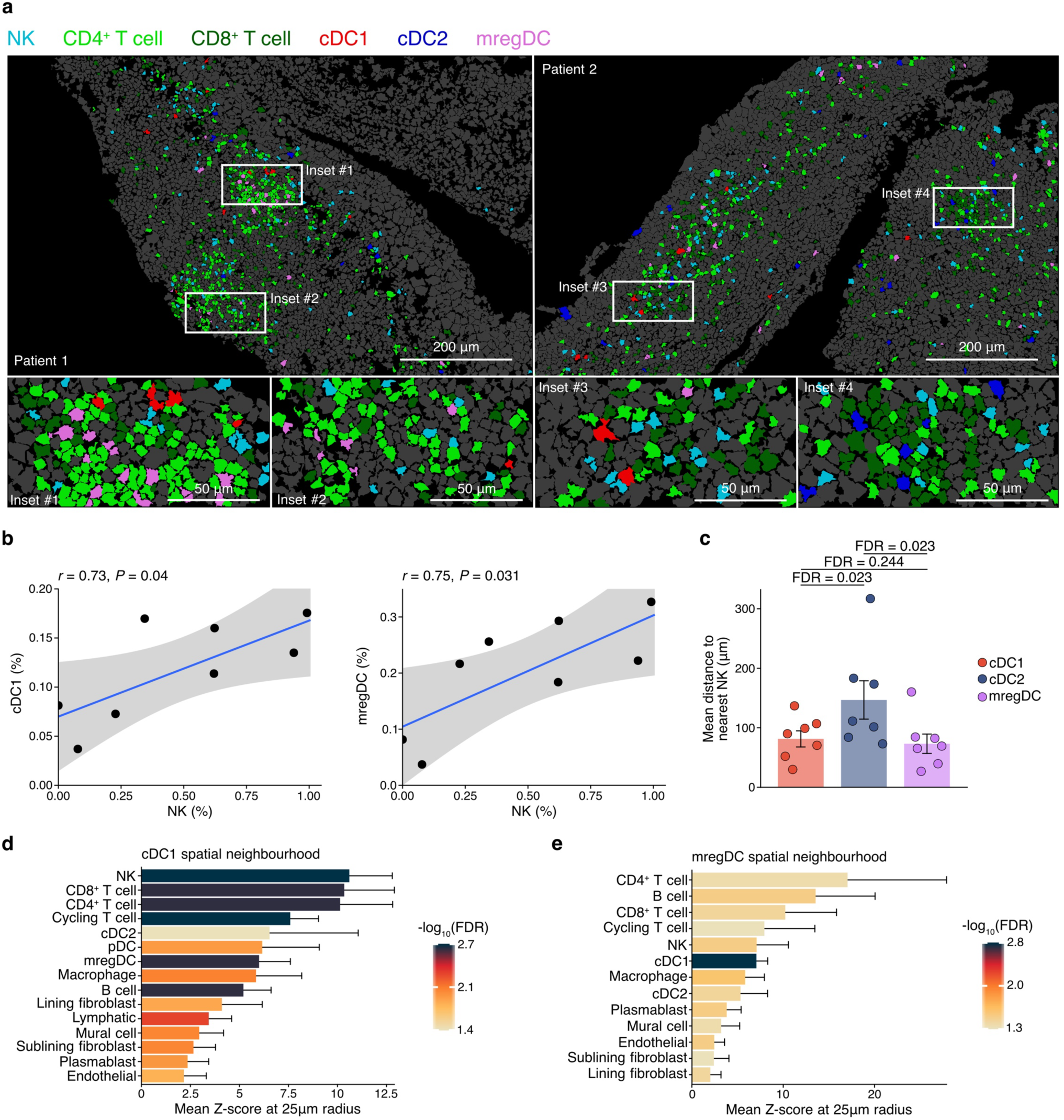
Synovial NK cells and cDC1s colocalise in distinct, T cell-enriched multicellular niches. **a**-**f**, Re-analysis of published Xenium spatial transcriptomics data derived from JIA synovial tissue^33^ (*n* = 8 patients). **a**, Representative cell segmentations of synovial tissue from two patients. The main panels display large tissue fields, while white bounding boxes indicate the magnified regions detailed in the corresponding bottom insets. Cells are coloured according to annotated subset. **b**, Pearson correlations of NK cell abundances with cDC1 (left) or mregDC (right) abundances, expressed as a percentage of all segmented cells. Points denote patient means calculated from tissue replicates. The solid line denotes linear regression, and the shaded region indicates the 95% CI. **c**, Mean spatial distances from cDC1, cDC2, and mregDC subsets to the nearest NK cell. **d**, **e**, Mean Z-scores demonstrating cell type enrichment within a 25 µm radius of cDC1s (**d**) or mregDCs (**e**). Bars indicate the mean ± s.d. Statistical significance was determined using paired two-tailed *t*-tests with Benjamini-Hochberg correction (**c**) and a one-sample *t*-test across patients against a null value of 0 (**d**, **e**), with error bars representing the mean ± 95% CI.

Single-cell distance metrics and spatial neighbourhood permutation analyses revealed that NK cells localised significantly closer to canonical cDC1s and mregDCs than to cDC2s (**Fig. 5c-e** and **Extended Data Fig. 5e**). These discrete NK cell-associated domains were densely populated by CD4^+^ and CD8^+^ T cells, forming spatially coordinated, T cell-enriched effector niches (**Fig. 5a, d, e**). Meanwhile, cDC2s occupied distinct tissue compartments and associated with T cells independently of NK cells and cDC1s (**Extended Data Fig. 5e**). This spatial segregation supports a model in which synovial NK cells selectively recruit cDC1s and promote the formation of localised T cell-cDC1 interaction niches within the inflamed joint.

### Cross-disease profiling identifies the pathogenic NK-cDC1 axis as a shared hallmark of inflammatory arthritis

To determine whether the XCL1-mediated NK-cDC1 axis is a disease-specific feature of JIA or a convergent inflammatory module across chronic arthropathies, we integrated previously published single-cell transcriptomic datasets from adult rheumatoid arthritis (RA)^49^, psoriatic arthritis^50^, and ankylosing spondylitis^51^. This unified atlas (**Extended Data Fig. 6a**) harmonised the NK and DC lineages across diverse clinical diagnoses and ages of onset.

A cross-disease meta-analysis revealed a striking conservation of the innate immune landscape. Irrespective of clinical phenotype, synovial fluid NK cells mapped to analogous transcriptional states identified in JIA: the infiltrating (C1), tissue-resident (C2), and activated (C3) subsets (**Fig. 6a and Extended Data Fig. 6b**). This conservation extended to their chemotactic programming, as synovial NK cells universally maintained high and selective expression of *XCL1* across these patient cohorts (**Fig. 6b**), coinciding with the profound enrichment of synovial fluid cDC1s exhibiting high *XCR1* expression (**Fig. 6c and Extended Data Fig. 6c**).

**Fig. 6:**
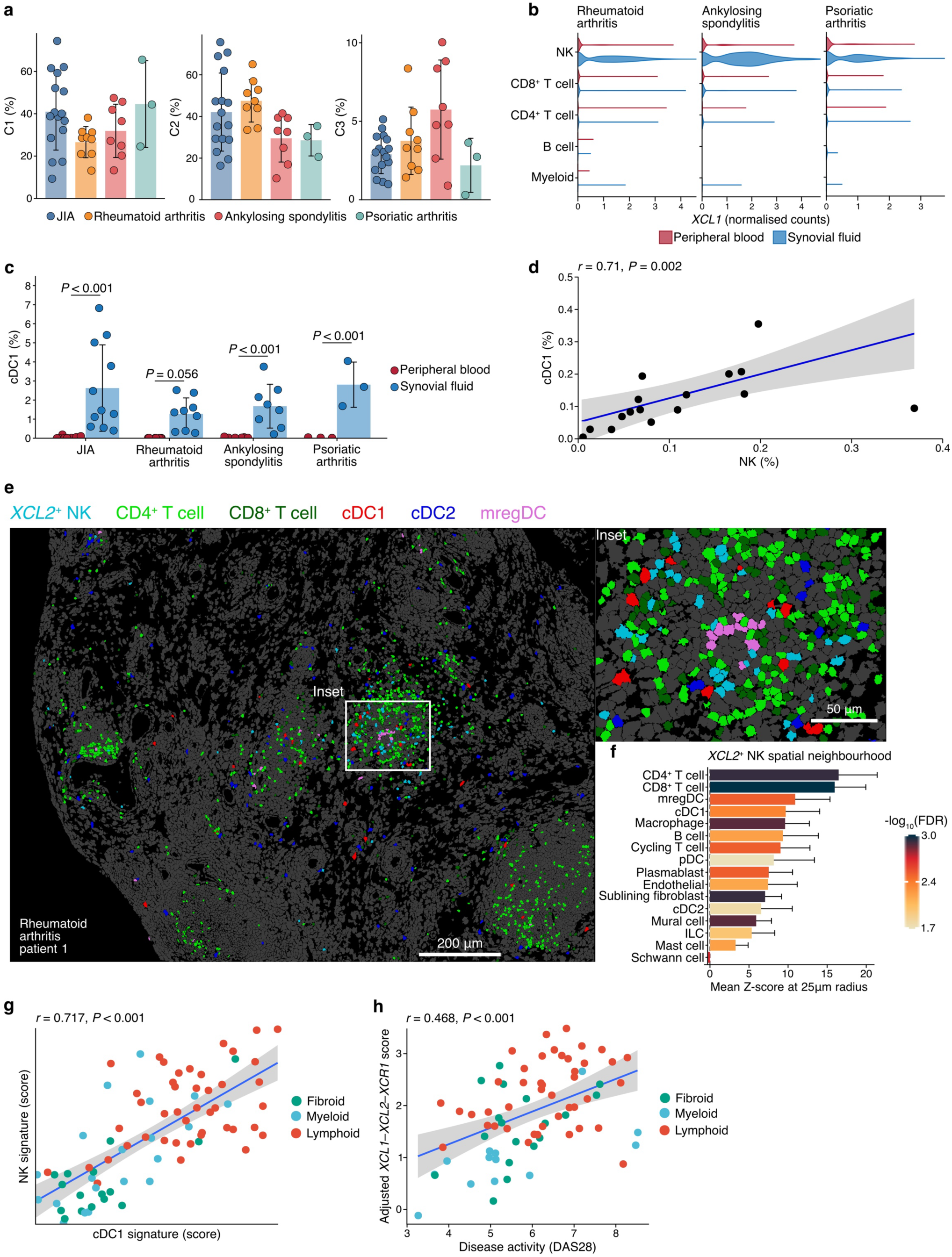
The XCL1/2-XCR1 immune axis is transcriptionally and spatially conserved across diverse forms of arthritis. **a**-**c**, Multi-cohort mononuclear CITE-seq analysis comparing blood and synovial fluid cells from JIA^33^ (paired, *n* = 16), rheumatoid arthritis^49^ (RA; unpaired, *n* = 9), ankylosing spondylitis^51^ (paired, *n* = 8), and psoriatic arthritis^50^ (paired, *n* = 3). **a**, Relative frequencies of NK cell subsets within total annotated synovial fluid NK cell populations. **b**, *XCL1* expression in annotated blood and synovial fluid immune cells across disease cohorts. **c**, Proportions of cDC1s, calculated as a percentage of all annotated cells, in blood and synovial fluid across disease cohorts. **d**-**f**, Re-analysis of published Xenium spatial transcriptomics of treatment-naïve RA synovial tissue^52^ (*n* = 17 patients), detailing the Pearson correlation between NK cell and cDC1 frequencies (**d**; points represent patient means from tissue replicates), representative tissue segmentations featuring a large tissue field and magnified inset (white box) (**e**), and mean Z-scores indicating cell type enrichment within a 25 µm radius of *XCL2*^+^ NK cells (**f**). **g, h,** Analysis of bulk synovial RNA-sequencing from the PEAC cohort^53^ (*n* = 82), demonstrating the correlation between NK cell and cDC1 modules (**g**) and the association between a composite *XCL1*, *XCL2*, and *XCR1* axis score and DAS28 macroscopic disease activity (**h**). In panels **d**, **g**, and **h**, solid lines depict linear regression, with shaded regions marking the 95% CI. Bars illustrate the mean ± s.d. Statistical significance was established using unpaired two-tailed *t*-tests with Benjamini-Hochberg correction (**a**), two-tailed *t*-tests or Mann-Whitney *U* tests contingent on data normality (**c**), and a one-sample *t*-test across patients (**f**).

To determine whether these convergent transcriptional signatures manifest as a conserved spatial architecture in situ, we re-analysed spatial transcriptomics of treatment-naïve RA synovial tissue^52^. Consistent with our findings in JIA, NK cell abundance significantly correlated with that of both cDC1s and mregDCs (**Fig. 6d and Extended Data Fig. 6d**), but not cDC2s (**Extended Data Fig. 6e**). Using *XCL2* as an available spatial probe, we distinguished *XCL2*^-^ and *XCL2*^+^ NK cell states, with unbiased spatial neighbourhood analysis revealing distinct spatial topologies for these subsets. Whereas *XCL2*^-^ NK cells associated primarily with the vasculature (**Extended Data Fig. 6f**), *XCL2*^+^ NK cells preferentially colocalised within densely occupied T cell-enriched niches alongside cDC1s and mregDCs (**Fig. 6e, f**).

Finally, to determine whether these localised associations extend to clinical disease activity, we analysed bulk synovial RNA-sequencing data from the Pathogenesis of Early Arthritis Cohort (PEAC)^53^. Transcriptomic module scoring confirmed that the concerted enrichment of NK cell and cDC1 signatures was highly characteristic of the densely infiltrated lymphoid and myeloid histological pathotypes (**Fig. 6g**). Furthermore, the aggregated expression of *XCL1*, *XCL2*, and *XCR1* was positively associated with systemic disease activity (DAS28) (**Fig. 6h**), independent of patient age, sex, disease duration and rheumatoid factor (RF) antibody positivity (β = 0.314, *P* < 0.001).

Collectively, these cross-cohort data demonstrate that the XCL1/2-XCR1 chemotactic axis is a conserved feature of inflammatory arthropathies, linking local microanatomical cellular coordination directly to macroscopic tissue pathology and systemic disease severity.

## Discussion

This study presents a comprehensive profile of blood and synovial innate lymphoid cells in treatment-naïve JIA patients. While single-cell transcriptomic approaches have begun to unveil NK cell heterogeneity at the periphery^20,24^, their transcriptional identities at sites of autoimmune inflammation remain largely unexplored. Our profiling reveals an innate lymphoid compartment dominated by the NK lineage, identifying transcriptional states that transcend the classic blood CD56^bright^ and CD56^dim^ dichotomy. We describe an infiltrating CD56^bright^-like subset that aligns with recent evidence demonstrating that circulating CD56^bright^ NK cells continuously infiltrate and adapt to solid tissues^25^. This population exists alongside CD103^+^ NK cells, which exhibit a tissue-residence signature similar to populations previously described in liver, uterus and small intestine^22,24,25^. Captured at the onset of disease, our data provide a unique snapshot of early tissue inflammation. We demonstrate that proliferating synovial NK cells exhibit a strongly activated expression signature, marked by the upregulation of cytokines such as TNF, IFN-γ, and GM-CSF, all of which can modulate autoimmune pathology in vivo^54–58^. Functionally, synovial NK cells are dominated by a chemokine-centred state, chiefly characterised by the spontaneous and highly specific expression of the paralogous chemokines XCL1 and XCL2. Although our protein-level validations focused exclusively on XCL1 due to the tight transcriptional co-expression and their shared cognate receptor (XCR1), both ligands likely act synergistically in vivo. Previous studies have demonstrated that NK cells are critical for the recruitment and spatial retention of cDC1s within tumour microenvironments^26,59^. The predominant expression of XCL1 in activated synovial NK cells, coupled with the elevated and selective expression of XCR1, rather than *CCR5*, on synovial cDC1s, indicates that the XCL1-XCR1 axis actively directs cDC1 recruitment to the inflamed joint. This specific interaction provides a mechanistic basis for the tight spatial colocalisation of cDC1s and NK cells observed in situ.

Previous single-cell studies in inflammatory arthritis have primarily focused on the characterisation of synovial cDC2s, as they are significantly more abundant than cDC1s in both blood and tissue^11,33^. However, our data demonstrate that the frequency of cDC1s, but not cDC2s, is associated with macroscopic disease pathology in JIA, consistent with findings in adult arthritis and tumour microenvironments^26,59^. A defining feature of canonical cDC1s is their capacity to cross-present antigen to CD8⁺ T cells, a process requiring CD4⁺ T cell-mediated licensing^60^. Memory CD8⁺ T cell responses can also be initiated within peripheral tissues through a tripartite interaction involving CD4⁺ T cells and recruited dendritic cells^61^. Such "triads" have been shown to play a critical role in antitumour immunity^62^. Our spatial analysis demonstrates that analogous multicellular niches emerge during autoimmune joint inflammation, where CD4⁺ and CD8⁺ T cells preferentially colocalise with cDC1s and NK cells. While memory CD8⁺ T cells might represent an adaptive source of synovial *XCL1* (ref ^39^), we propose that rapid, antigen-independent XCL1 production by locally activated NK cells shapes the initial assembly of these early niches. By spatially anchoring cDC1s within T cell-rich zones, this early NK-derived gradient facilitates the formation of cross-presentation-competent triads, establishing autonomous effector loops that sustain chronic inflammation across affected joints.

Within these dense niches, the local microenvironment also drives extensive functional adaptation. Although previous profiling of the adult RA synovium mapped the *CCR7*⁺ *LAMP3*⁺ mregDC state exclusively to the cDC2 lineage^11^, our data demonstrate that synovial cDC1s also converge into this shared, highly conserved state^44^, downregulating *WDFY4* in the process. This reveals a bifurcated functional fate for the recruited cDC1 pool. While a large fraction of canonical cDC1s retains *WDFY4* expression, thereby remaining competent for cross-presentation, a subset phenotypically transitions into mregDCs. Although losing *WDFY4* signifies a departure from this canonical function^48^, cDC1-derived mregDCs uniquely upregulate *IL12B* (IL-12p40). This may set up a self-amplifying positive feedback loop: NK cells recruit XCR1^+^ cDC1s via XCL1, enabling their local maturation into IL-12-producing mregDCs that, in turn, boost local NK cell and T_h_1 responses. Because mregDCs highly express regulatory checkpoints such as PD-L1 and PD-L2, likely functioning as homeostatic brakes, their net role in arthritis pathogenesis warrants careful evaluation.

While our multi-modal human data strongly suggest these cellular dynamics, definitive proof of real-time in situ mregDC function and triad formation will rely on future in vivo lineage-tracing models. Nevertheless, the identification of a conserved NK-cDC1 axis across chronic arthropathies inherently provides a compelling rationale for early therapeutic intervention. Because this circuit is already established in treatment-naïve patients, it likely operates as a fundamental upstream driver of joint inflammation. Indeed, the in vivo neutralisation of XCL1 ameliorates models of arthritis^39,63^, underscoring its unique pathogenic contribution and targetability. Disrupting these upstream innate networks offers a promising, disease-modifying strategy to dismantle pathogenic microanatomical niches universally active in chronic synovial inflammation.

## Methods

### Human participants

Children diagnosed with non-systemic onset juvenile idiopathic arthritis (JIA) were recruited from the Department of Pediatric Respiratory Medicine, Immunology and Critical Care Medicine at Charité - Universitätsmedizin Berlin between October 2020 and August 2025. We obtained written informed consent from all participants or their legal guardians prior to sample collection. The study adhered to the ethical principles of the Declaration of Helsinki and was approved by the local ethics committee of Charité Berlin (EA2/113/20, EA2/048/26). We obtained healthy control blood from buffy coats supplied by the Deutsches Rotes Kreuz Blutspendedienst Nord-Ost, with approval from the Charité Berlin ethics committee (EA1/243/24). Patient-associated metadata are provided in Supplementary Table 1.

### Clinical sample collection and mononuclear cell isolation

Whole blood from patients and healthy donors was collected in Vacuette tubes (Greiner Bio-One, 455084). To isolate serum, blood was collected in Vacutainer SST tubes (BD, 367986), centrifuged, aliquoted, and stored at −20°C until analysis. We obtained synovial fluid via syringe aspiration. If aspirate volumes were low, we pooled aspirates from multiple joints of the same patient. We recorded the initial mass and final dilution factor in PBS containing 0.1% bovine serum albumin (BSA) to normalise downstream measurements.

We isolated peripheral blood mononuclear cells (PBMCs) by diluting whole blood 1:1 with PBS, followed by density gradient centrifugation using Ficoll (GE Healthcare, GE17-1440-02). Synovial fluid mononuclear cells (SFMCs) were kept on ice and isolated by passing aspirates directly through 70 µm cell strainers (Miltenyi, 130-110-916) and washing with PBS/BSA. Isolated PBMCs and SFMCs were either used immediately or cryopreserved in foetal bovine serum (FBS) supplemented with 10% dimethyl sulfoxide.

### Ex vivo cell stimulation and cytokine quantification

Cryopreserved PBMCs and SFMCs were thawed in complete RPMI 1640 medium (Gibco, 11875093) supplemented with 10% FBS (Gibco, A5256701), 100 U/mL penicillin-streptomycin (Gibco, 15140122), and 25 U/mL Benzonase (Sigma-Aldrich, E1014). Cells were washed and cultured in this complete medium at 37°C in a 5% CO_2_ atmosphere under specific stimulation conditions. To assess intracellular cytokines, we either cultured cells immediately with GolgiStop (monensin, 0.6 μL/mL, BD, 554724) and GolgiPlug (brefeldin A, 1 μL/mL, BD, 555029) or stimulated them for 2 hours before adding these transport inhibitors for a further 4 hours.

For receptor cross-linking, we conjugated 30 μg of biotinylated target antibody to 500 μL of anti-biotin MACSi beads (Miltenyi, 130-092-357). The cross-linking panel comprised anti-CD16 (clone 3G8, 302004) alongside an equimolar mixture of antibodies targeting CD2 (RPA-2.10, 100103), 2B4 (eBioC1.7, 329504), NKp46 (9E2, 331906), NKp30 (P30-15, 325206), NKG2D (1D11, 320804), and DNAM-1 (11A8, 338326), all sourced from BioLegend. Antibody-loaded beads were added to cell cultures at a bead-to-cell ratio of 5:1 to ensure saturating cross-linking across the multiplexed receptor panel. We performed cytokine stimulations with IL-12 (10 ng/mL, Miltenyi, 130-096-704), IL-15 (10 ng/mL, Miltenyi, 130-095-760), or IL-18 (100 ng/mL, MBL International, B001-5). Positive controls were stimulated with phorbol myristate acetate (20 ng/mL, Sigma-Aldrich, P8139) and ionomycin (1 μg/mL, Sigma-Aldrich, I0634).

### Enzyme-linked immunosorbent assay

Patient serum and synovial fluid supernatant samples were thawed simultaneously. We determined XCL1 protein concentrations using the Human XCL1 ELISA Kit (Abcam, ab264620) according to the manufacturer’s protocols. All samples were analysed in duplicate using a Spark Multimode Microplate Reader (TECAN, 30086376). Dilution factors for synovial fluid supernatant measurements were corrected for by multiplying the initial aspirate dilution during SFMC isolation. One patient serum measurement fell below the manufacturer’s specified sensitivity threshold (28.3 pg/mL) and was excluded from subsequent analyses.

### Flow cytometry and cell sorting

Cells were incubated with a fixable viability dye at room temperature in PBS for 10 minutes, washed, and stained with extracellular antibodies in PBS/BSA. Intracellular staining was performed using the eBioscience Foxp3 / Transcription Factor Staining Buffer Set (Thermo Fisher Scientific, 00-5523-00). Data were acquired on a five-laser Cytek Aurora full-spectrum cytometer and analysed using FlowJo software (v10.11, BD Biosciences). For analytical flow cytometry, specific populations were identified following doublet exclusion and live cell gating.

For single-cell sequencing, live cells were sorted using a FACSAria II (BD Biosciences). Innate lymphoid cells were sorted from fresh PBMCs and SFMCs as viable, CD14^-^ CD15^-^ CD123^-^ CD235a^-^CD19^-^ CD3^-^ CD7^+^ events and pooled at a 1:1 ratio. Dendritic cells (DCs) were sorted from thawed samples as CD45^+^ CD19^-^ CD235a^-^ CD3^-^ CD88^-^ events. These were further subdivided into plasmacytoid DCs (CD303^+^), type 2 conventional DCs (CD11c^+^ CD1c^+^), and type 1 conventional DCs (CD11c^+^ CD141^+^). Before sorting, cells were labelled with TotalSeq-A or TotalSeq-C antibodies for antibody-derived tag (ADT) recovery, alongside specific hashtag oligonucleotides (HTOs) per patient, and per origin, for demultiplexing. Following sorting, DC subsets were pooled to ensure equal representation. Lists of antibodies used for flow cytometry, cell sorting, and CITE-seq are provided in Supplementary Table 2.

### Single-cell RNA library preparation and sequencing

Sequencing libraries were prepared targeting 25,000 cells per reaction. Innate lymphoid cell libraries were generated using the Chromium Next GEM Single Cell 5’ v2 kit with feature barcoding (10x Genomics, CG000330). DC and total immune cell libraries were generated using the Chromium Next GEM Single Cell 3’ v3.1 kit (10x Genomics, CG000204), modified for ADT and HTO recovery. We amplified ADT and HTO libraries using KAPA HiFi ReadyMix (Roche, 07958927001) with sample-specific indices, followed by SPRIselect purification (Beckman Coulter, B23319). Library quality was confirmed using a Fragment Analyzer (Advanced Analytical) and Qubit 2.0 (Thermo Fisher Scientific). Sequencing was performed on an Illumina NovaSeq 6000 instrument with a 1% PhiX spike-in.

### Single-cell RNA sequencing preprocessing and quality control

We demultiplexed base call files using bcl2fastq (v2.20.0.422, Illumina) and the output was mapped to an optimised GRCh38 reference genome^64^. Ambient RNA and ADT/HTO background was then corrected using CellBender^65^ (v0.3.2). To facilitate patient demultiplexing, we genotyped exonic variants using cellsnp-lite^66^ (v1.2.4) and vireo^67^ (v0.2.3). We then imported the corrected expression matrices into R^68^ (v4.3.3) as Seurat (v5.4.0) objects, subsequently removing pseudogenes and long non-coding RNAs.

We applied quality control to exclude cells exhibiting outlier distributions of mitochondrial, ribosomal, and haemoglobin transcripts per library. We further excluded potential platelet contaminants (*PPBP* expression > 0.001%) and applied heat shock protein thresholds (retaining cells with > 0.01% and ≤ 2.5% HSP family gene expression). Surface HTO and ADT data underwent per-feature and per-cell centred log-ratio (CLR) transformation, respectively. We identified and removed doublets using vireo and DoubletFinder^69^ (v2.0.4). For DoubletFinder, we adjusted the expected doublet rate per library using the modelHomotypic function on Leiden clusters (resolution 0.1). Optimal pK values were computationally determined per library using the paramSweep function, maintaining pN = 0.25 across 30 principal components.

### Integration, clustering, and cell type annotation

We normalised gene expression data using depth-relative proportional fitting^70^ prior to integrating data across patients using Harmony^71^ (v2.1.1). To annotate cell types, we utilised Azimuth^72^ (v0.5.0) reference mapping alongside literature-curated markers, followed by Leiden^73^ (v0.4.3.1) clustering. For individual objects, we applied an initial clustering resolution of 0.5 to identify and exclude non-target clusters. Finally, we reintegrated and subclustered the remaining cells at resolutions of 0.3 and 0.1 to obtain high-resolution final subsets.

### Differential gene expression and pathway enrichment

We performed differential gene expression analysis using two approaches. For multiple comparisons between clusters, we used the Seurat FindAllMarkers function with a two-sided Wilcoxon rank-sum test, restricting the analysis to genes expressed in at least 10% of cells. We defined significance as a log_2_ fold change ≥ 0.5 and a Bonferroni-adjusted P value ≤ 0.001. We removed mitochondrial, ribosomal, and *MALAT1* transcripts from the outputs post-computation. To mitigate pseudoreplication artefacts during cross-tissue and distinct cell population pairwise comparisons, we generated pseudobulk aggregated count matrices using the Seurat AggregateExpression function. We performed differential expression analysis on these matrices using DESeq2^74^ (v1.42.1) coupled with apeglm^75^ (v1.24.0) effect size shrinkage. Prior to model fitting, we filtered the dataset to retain only genes demonstrating 10 or more counts across a minimum of three independent pseudobulk samples. For cross-tissue analyses, we implemented a paired design to control for patient-specific variance. We set significance thresholds for pseudobulk analyses at an absolute log_2_ fold change ≥ 0.5 and a false discovery rate (FDR) ≤ 0.001, excluding mitochondrial and ribosomal transcripts. All differential expression results supporting the figures and extended data are provided in Supplementary Table 3.

We calculated Gene Ontology Biological Process enrichment using clusterProfiler^76^ (v4.10.1). Input gene lists required a log_2_ fold change ≥ 1.25 and an FDR ≤ 0.001. We determined enrichment significance using a hypergeometric test with Benjamini-Hochberg adjustment, setting the significance threshold at an FDR *q* value ≤ 0.05.

### Transcriptional module and cell cycle scoring

We computed transcriptional module scores using the Seurat AddModuleScore function. We defined the blood CD56^bright^ NK cell signature by comparing blood CD56^bright^ against CD56^dim^ NK populations. Cytokine-stimulated NK cell modules^40^ comprised genes significantly upregulated following three hours of in vitro stimulation with either IL-2 and IL-15, or IL-12 and IL-18. The acute T cell activation module comprised time-resolved consensus genes^41^ significantly upregulated one hour following in vitro anti-CD3 and anti-CD28 stimulation, retaining only transcripts verified against unstimulated negative controls. To predict the lineage origin of mregDCs, we defined cDC1 and cDC2 signatures as the intersection of subset-defining genes conserved across both blood and synovial fluid (absolute log_2_ fold change ≥ 1, FDR ≤ 0.001). We assigned a specific lineage if a cell module score exceeded both the opposing module score and the standard deviation of the target module across the mregDC population.

We computed cell cycle scores using the Seurat CellCycleScoring function. We defined actively cycling cells using standard Seurat cell-cycle genes^77^. To quantify phenotypic shifts, we calculated the mean Euclidean distance in principal component space between dendritic cell subsets. We computed these distances per patient using variance-stabilising transformed counts of differentially expressed genes (absolute log_2_ fold change ≥ 0.5, FDR ≤ 0.001).

To project these high-resolution states clinically, paired bulk synovial RNA-sequencing data from the PEAC cohort^53^ were quantified using kallisto^78^ (v0.46.1) against the Ensembl GRCh38 115 cDNA reference, log_2_(abundance + 1)-transformed, and scored for core NK and cDC1 derived signatures using GSVA^79^ (v1.50.5). To evaluate clinical associations, the *XCL1-XCR1* axis score (mean transcript Z-score) was correlated with DAS28 using ordinary least squares and robust linear regression (bisquare weighting), adjusting for age, sex, disease duration, and rheumatoid factor status. Model stability was validated via leave-one-gene-out sensitivity analysis and 2,000-iteration bootstrapping.

### Trajectory inference

For trajectory inference, we isolated canonical cDC1 and cDC2 lineages alongside synovial fluid mregDCs. To mitigate sequencing chemistry batch effects, we reintegrated the lineages independently using canonical correlation analysis^80^. We defined developmental progression stages a priori based on tissue origin and cell type annotations. We inferred pseudotime by applying Slingshot^81^ (v2.7.0) to the reduced dimensions, using predefined stages as cluster labels without specifying a root or terminal cell state. We then normalised pseudotime values to a range of 0 to 1, visualising relative gene expression dynamics along the inferred trajectory using generalised additive model fits, indicating 95% confidence intervals with shaded areas.

### Spatial transcriptomics analysis

We obtained Xenium spatial data from previously published cohorts^33,52^. We refined initial cell segmentations using baysor^82^ (v0.71) with a 10-molecule minimum threshold and a prior segmentation confidence of 0.7. We quantified spatial colocalisation using spatstat^83^ (v3.5-1) through a hierarchical permutation framework. Toroidal shift permutations (*n* = 1,000) generated Z-scores at several radii to represent observed versus expected nearest neighbours. We assessed cross-cohort enrichment using a one-sample t-test against a null hypothesis mean Z-score of 0, applying Benjamini-Hochberg FDR correction (Supplementary Table 3). To quantify spatial proximity between specific cell populations, we computed minimum two-dimensional Euclidean distances from target cells to the nearest focal cell type independently within each tissue section. This was achieved using a k-d tree nearest-neighbour algorithm implemented in RANN^84^ (v2.6.2).

### Public dataset integration

We incorporated publicly available single-cell RNA-sequencing data from juvenile idiopathic arthritis^33^, rheumatoid arthritis^49^, ankylosing spondylitis^51^, and psoriatic arthritis^50^ cohorts. We realigned these datasets to the optimised GRCh38 reference genome^64^, corrected background noise using CellBender^65^, and integrated all datasets by patient using Harmony.

### Statistical analysis

We performed statistical calculations in R^68^ (v4.3.3). We evaluated relationships between donor-averaged gene expression profiles and between spatial or tissue-specific cell proportions using Pearson correlation tests (cor.test). We modelled and visualised these relationships using linear regressions (lm). We selected pairwise comparison methods based on Shapiro-Wilk normality testing. For fully paired data, we used a paired *t*-test or Wilcoxon signed-rank test; for unpaired data, we used an unpaired *t*-test or Mann-Whitney U test. We analysed groups with partial patient overlap (three or more shared patients) using linear mixed-effects models with the patient included as a random effect (lme4^85^ (v 1.1-38) and lmerTest^86^ (v3.2-0); Satterthwaite degrees of freedom). Groups with fewer than three shared patients were analysed as unpaired. We corrected for multiple comparisons using the Benjamini-Hochberg method; single comparisons remained uncorrected. All statistical tests required a minimum of three observations per group.

## Supporting information

Supplementary Table 1: Patient demographics

Supplementary Table 2: Antibodies

Supplementary Table 3: Differentially expressed genes

## Data availability

Raw FASTQ files will be accessible through the German Human Genome-Phenome Archive (GHGA; [accession placeholder]) upon publication. Access to these controlled data requires the submission of a formal project application and subsequent approval by the designated Data Access Committee (DAC).

Processed, non-identifiable matrices (CellBender outputs and Seurat objects) are deposited on Zenodo under accession no. 18514472.

To facilitate interactive exploration, visualisation of gene and antibody expression for the innate lymphoid cell, dendritic cell, and mononuclear cell objects is available on the CELLxGENE platform (cellxgene.cziscience.com/collections/10eb236d-d42d-45b8-8363-c2dcf865f388).

## Code availability

All code to reproduce the analyses performed in this paper will be made available upon publication on GitHub at github.com/ollieeknight/paper_nk_cdc1_in_jia.

## Acknowledgements

We thank the patients and their legal guardians for their participation and the provision of clinical samples. We acknowledge the Flow Cytometry Core Facility at the Deutsches Rheuma-Forschungszentrum Berlin (T. Kaiser, J. Kirsch, and K. Heinrich) for assistance with cell sorting, and the High-Performance Computing for Research cluster at the Berlin Institute of Health for computational resources. We are grateful to members of the Romagnani laboratory, T.M. Campbell, and S.G. San-Miguel for critical reading of the manuscript. We also thank M. Klemm, L. Puccio, and C. Hobe for technical assistance, and M. Massoud, G. Guerra, and C.C. Goetzke for valuable support.

## Contributions

C.R., T.K. and M.F.M. established the initial research framework. O.C.K., T.R. and C.R. designed the study. T.K., A.S.L.v.S. and C.W. performed the clinical evaluations, recruited the patients and collected the paired blood and synovial fluid samples. O.C.K. directed the experimental work and data analysis, which included performing the single-cell sequencing experiments with T.R., conducting the stimulation assays with C.G. and undertaking all bioinformatics and computational analyses, with integration of metadata and analytical input provided by C.B, C.B.M and A.P.C. C.R. secured the primary funding for the study. O.C.K. and C.R. wrote the manuscript. All authors reviewed and edited the manuscript.

