## Extended Data Figures for "Tissue-adapted NK cells shape pathogenic cDC1 niches in early arthritis"

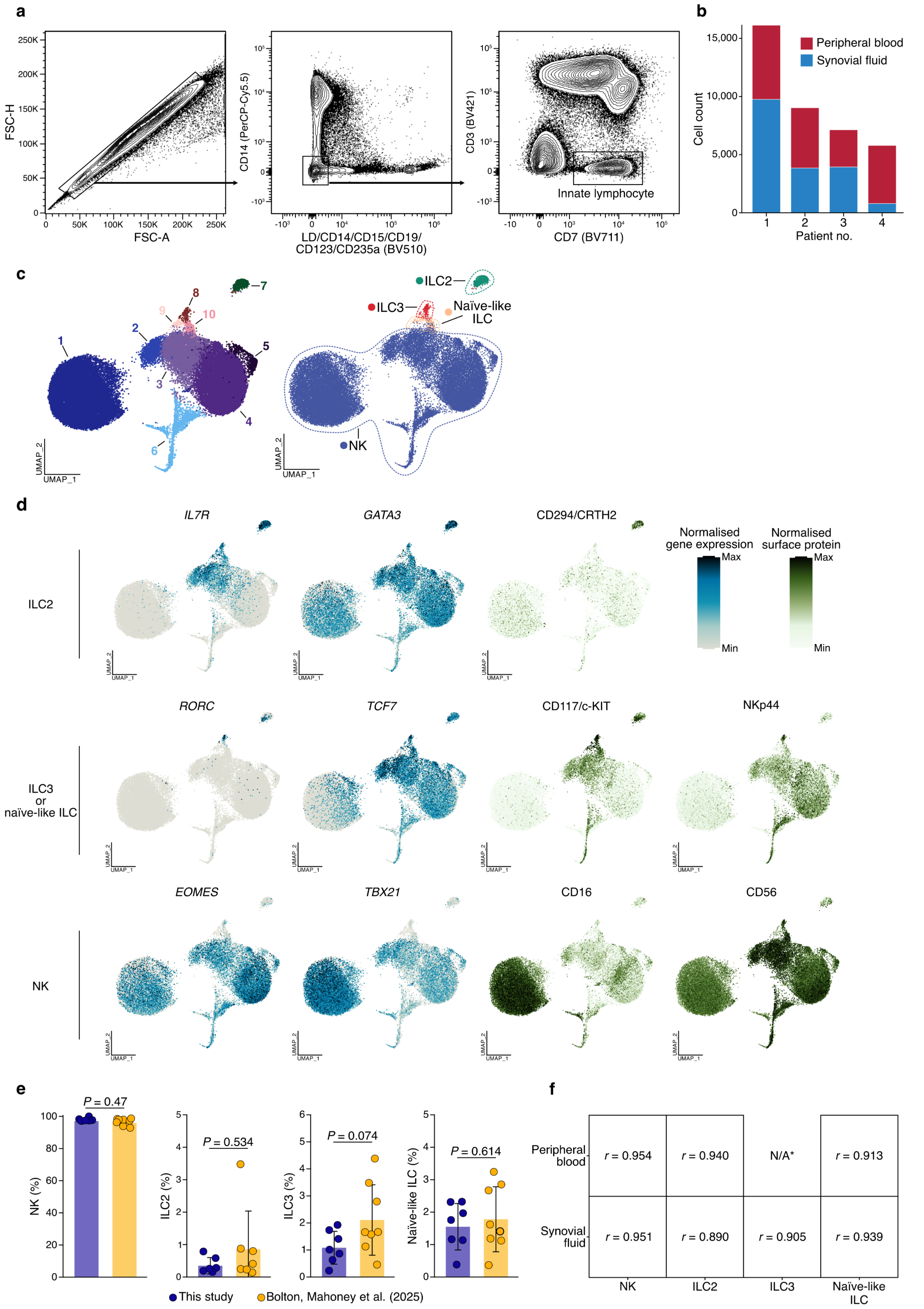


**Extended Data 1: Isolation and identification of innate lymphoid cell subsets by CITE-seq. a**, Sorting strategy applied for the isolation of ILCs prior to CITE-seq analysis. The panel displays representative flow cytometry gating for JIA synovial fluid. **b**, Total patient cell counts recovered by CITE-seq, delineated by patient and tissue origin. **c, d,** Transcriptional and surface protein definitions of the cells recovered by CITE-seq. Panels display integrated UMAPs coloured by ordered Leiden cluster or manual subset annotation (**c**), alongside feature plots (**d**) demonstrating the CITE-seq expression of lineage marker genes in blue and Antibody-Derived Tag (ADT) surface proteins in green. Panels represent specific subsets as follows: top, ILC2s (*IL7R*, *GATA3*, and CD294/CRTH2); middle, ILC3s or naïve-like ILCs (*RORC*, *TCF7*, CD117/c-KIT, and NKp44); and bottom, NK cells (*EOMES*, *TBX21*, CD16, and CD56). **e, f,** Integration with a published dataset[1] to compare the frequencies of NK and ILC subsets (**e**) and the Pearson correlation coefficients of their mean gene expression profiles across blood and synovial fluid origins (**f**). Bars denote the mean ± s.d. Statistical comparisons relied on an unpaired two-tailed *t*-test (**e**).

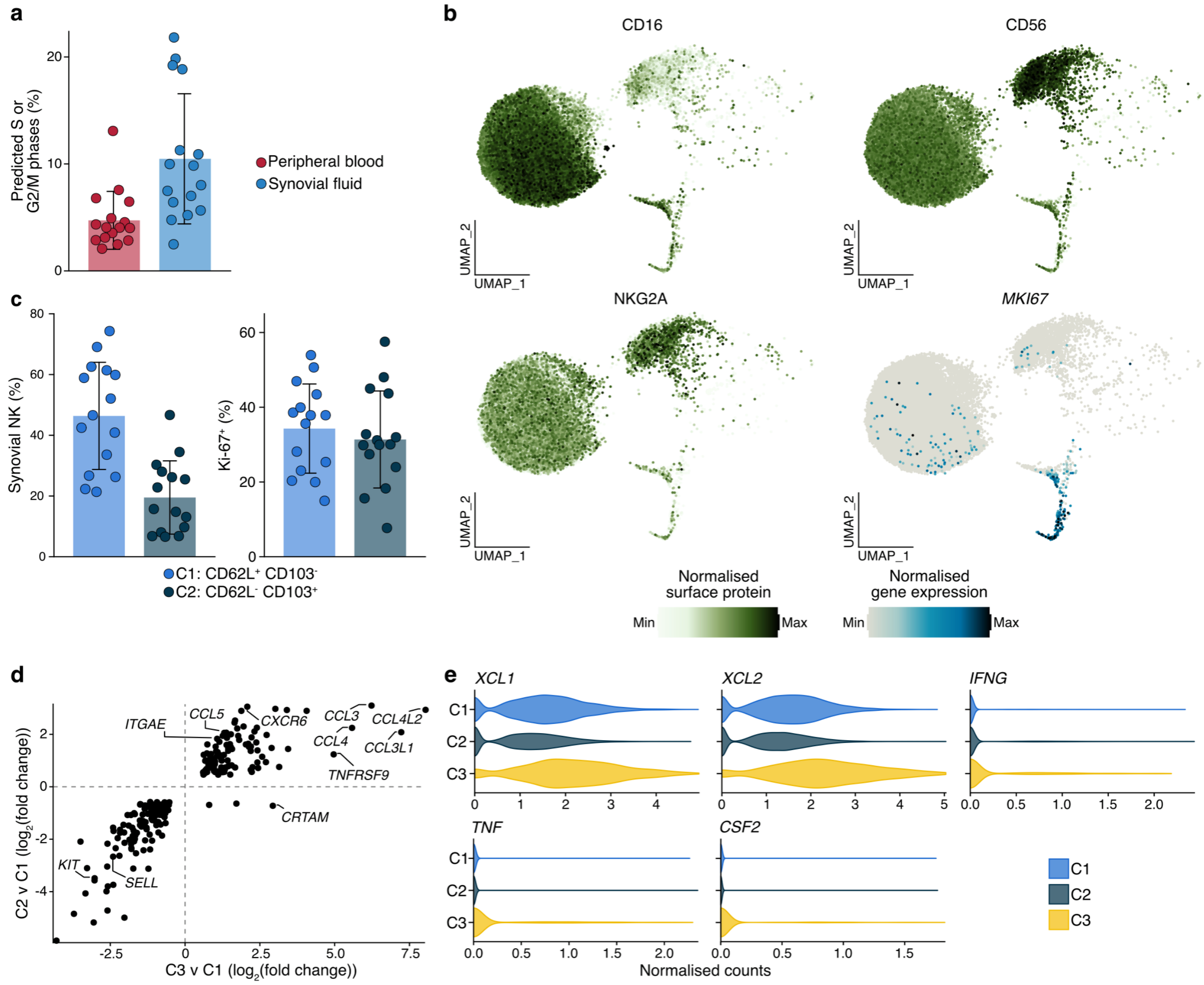


**Extended Data 2: Transcriptional and surface marker definitions of JIA NK cell states. a**, Predicted proportions of S and G2/M phase NK cells in blood and synovial fluid, calculated as a percentage of total annotated NK cells (CITE-seq). **b**, Feature plots illustrating the normalised expression of surface proteins CD16, CD56, and NKG2A, alongside *MKI67* in blood NK cells. **c**, Distribution of CD62L$^{+}$ CD103$^{-}$ and CD62L$^{-}$ CD103$^{-}$ populations among total synovial NK cells (left) and Ki-67$^{+}$ NK cells (right), evaluated by flow cytometry. **d**, Scatter plot comparing the $\log_2$ fold changes of differentially expressed genes in synovial fluid NK cells, capturing the intersection of pairwise C3 versus C1 and C2 versus C1 comparisons. **e**, Violin plots presenting the expression of *XCL1*, *XCL2*, *IFNG*, *TNF,* and *CSF2* (GM-CSF) across synovial fluid NK subpopulations. Statistical significance was determined using a paired Wald test via DESeq2, setting C1 as the reference (**d**).

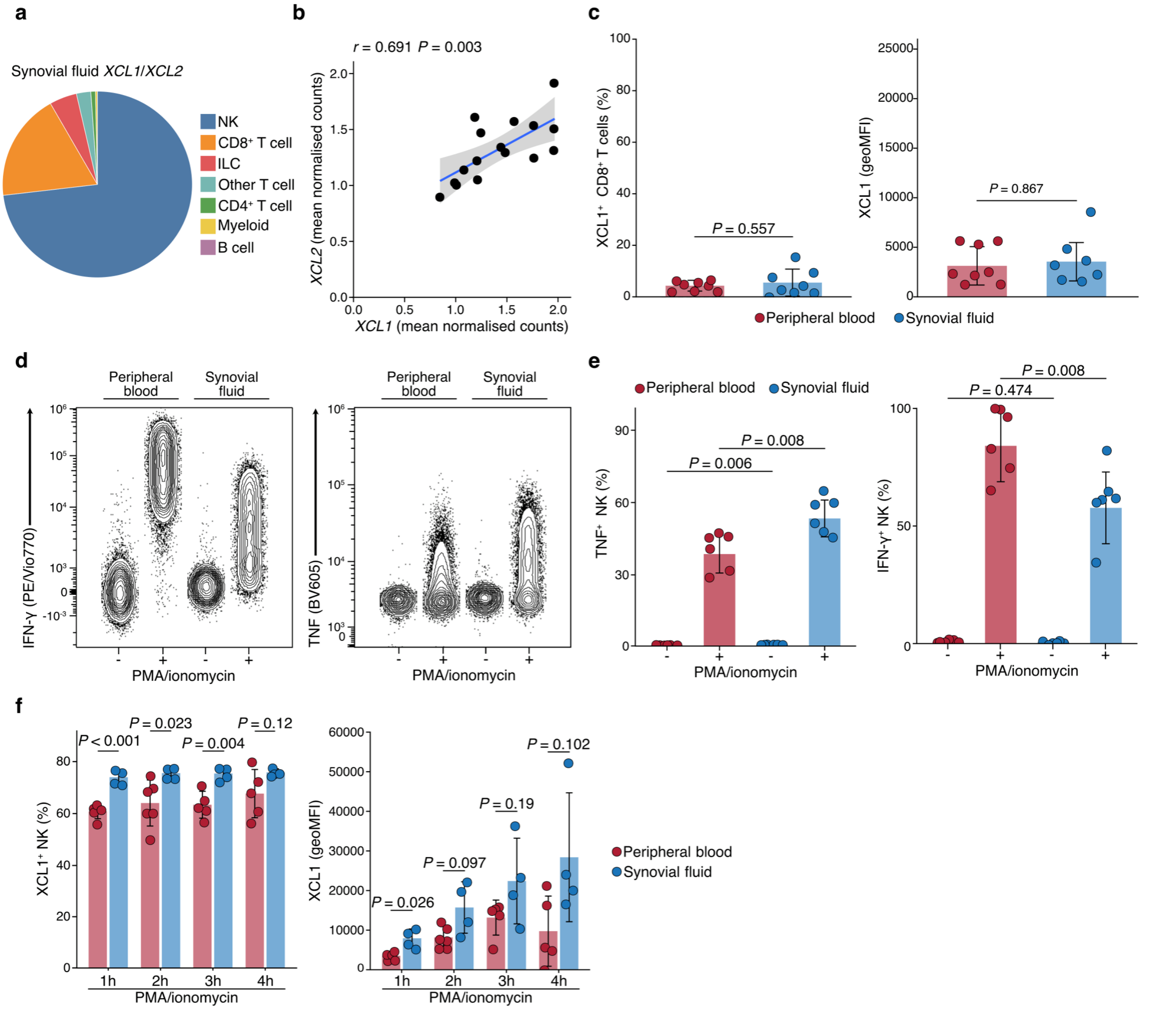


**Extended Data 3: Functional evaluation of XCL1 production by NK cells. a**, Cellular composition of the total *XCL1*- and *XCL2*-expressing pool within JIA synovial fluid (CITE-seq, *n* = 11). Proportions were calculated per patient prior to averaging. **b**, Pearson correlation of normalised *XCL1* and *XCL2* expression within synovial fluid NK cells (CITE-seq, *n* = 11 patients). The solid line delineates linear regression, with the shaded region indicating the 95% CI. **c**, Frequencies (left) and geoMFI (right) of $XCL1^+$ $CD8^+$ T cells in blood (*n* = 8) and synovial fluid (*n* = 7; 7 paired, 1 unpaired), calculated as a percentage of total $CD8^+$ T cells (gated as live, $Lin^-$ $CD3^+$ $CD8^+$ events). Flow cytometric analysis was performed after a six-hour unstimulated culture containing brefeldin A and monensin. **d-f,** Flow cytometric determination of intracellular cytokine production by blood and synovial fluid NK cells, following culture with or without PMA and ionomycin in the presence of brefeldin A and monensin. Panels display representative plots (**d**) and quantified frequencies (**e**) of $TNF^+$ and IFN-$\gamma^+$ NK cells after six hours of culture. These are presented alongside the frequencies and geoMFI of $XCL1^+$ NK cells, monitored across a four-hour stimulation time course (**f**; *n* = 5 for blood, *n* = 4 for synovial fluid). Bars signify the mean ± s.d. Statistical significance was calculated using paired *t*-tests or Mann-Whitney *U* tests, contingent upon data normality (**c**, **e**, **f**; restricted to paired observations).

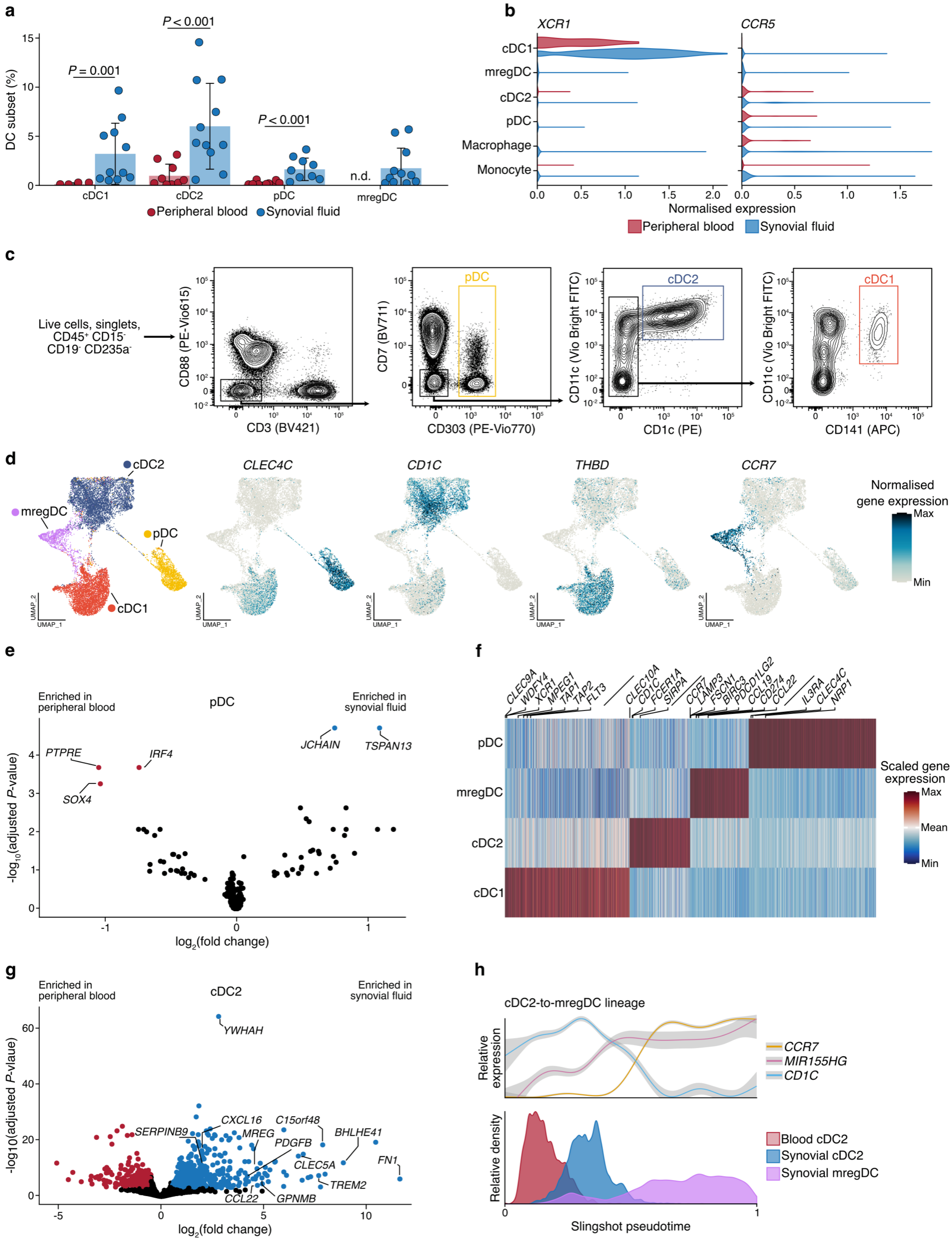


**Extended Data 4: Transcriptional profiling and functional signatures of synovial dendritic cells. a**, Abundance of DC subsets within the total annotated mononuclear cell populations of blood and synovial fluid (CITE-seq, $n$ = 11). **b**, Normalised *XCR1* and *CCR5* expression across annotated blood and synovial fluid myeloid cells (encompassing monocyte, macrophage, and DC populations) derived from CITE-seq data. **c**, Sorting strategy applied to isolate DC subsets prior to single-cell sequencing. The panel demonstrates a representative flow cytometry staining from JIA synovial fluid. **d**, Integrated UMAP of blood (1,863 cells) and synovial fluid (13,200 cells) DCs, coloured by subset (left). Accompanying feature plots (middle, right) map the gene expression of *CLEC4C* (CD303), *CD1C* (CD1c), *THBD* (CD141), and *CCR7* (CITE-seq). **e**, **g**, Differentially expressed genes between synovial fluid and blood pDCs (**e**) or cDC2s (**g**) generated from CITE-seq data. **f**, Heatmap detailing row-scaled differentially expressed genes that delineate DC subpopulations, with selected markers highlighted. **h**, Slingshot pseudotime trajectory modelling the cDC2-to-mregDC transition. The top panel demonstrates the relative normalised expression of lineage-specific and maturation genes along the pseudotime axis; lines show generalised additive model fits, with shaded regions marking the 95% CIs. The bottom panel indicates the relative density of cells across the inferred trajectory. Bars designate the mean ± s.d. Statistical significance was determined using paired two-tailed *t*-tests (**a**), Wald tests with Benjamini-Hochberg correction via DESeq2 (**e**, **g**), and a two-sided Wilcoxon rank-sum test with Bonferroni correction (**f**).

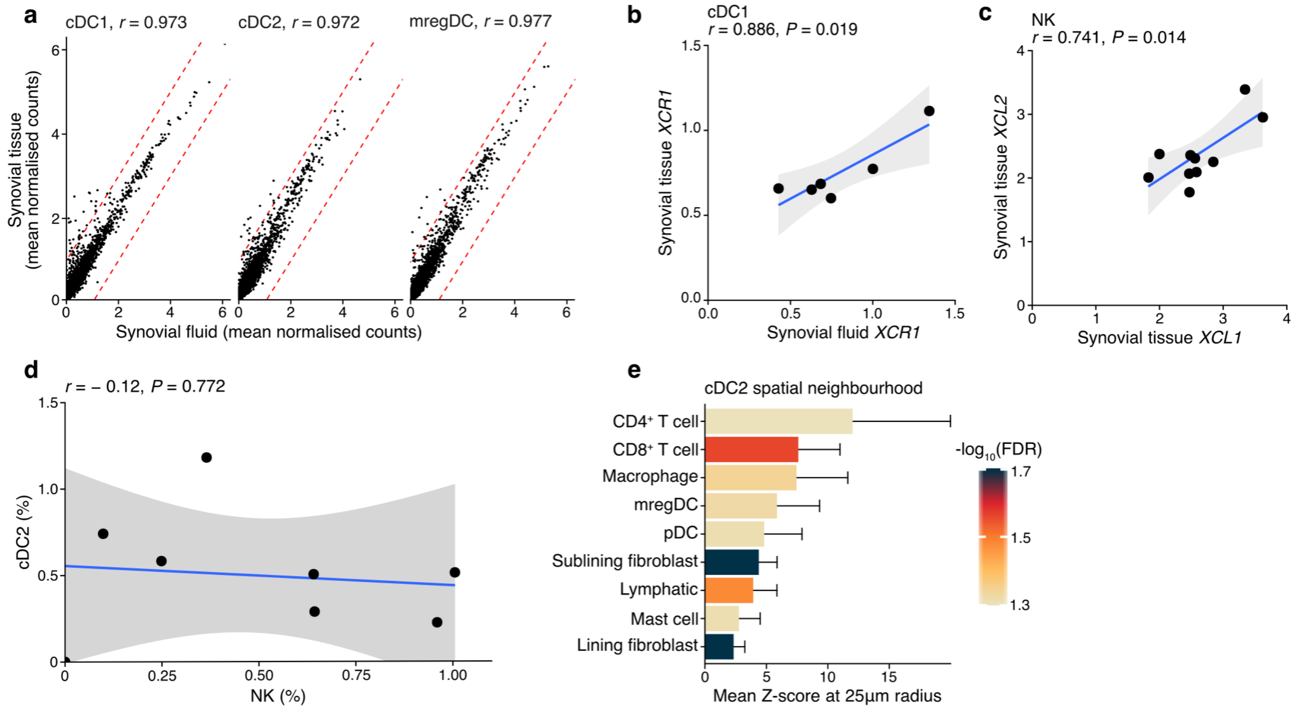


**Extended Data 5: Conservation of the XCL1/2-XCR1 axis and spatial segregation of cDC2s in the inflamed synovium.** **a**, Scatter plots and Pearson correlation coefficients of mean gene expression profiles for cDC1s, cDC2s, and mregDCs across synovial fluid and tissue, utilising a published CITE-seq cohort[1]. Dashed red lines signify a $\log_2$ fold change of 1 toward either tissue origin. **b**, Pearson correlation of *XCR1* expression in cDC1s sourced from paired synovial fluid and tissue (CITE-seq, $n$ = 6 patients). The solid line indicates linear regression, while the shaded region indicates the 95% CI. **c**, Pearson correlation of *XCL1* and *XCL2* expression in synovial tissue NK cells (CITE-seq, $n$ = 10). The solid line represents linear regression, with the shaded region indicating the 95% CI. **d**, **e**, Re-analysis of published Xenium spatial transcriptomics data from JIA synovial tissue[1] ($n$ = 8 patients). Panel **d** shows the correlation between NK cell and cDC2 abundances (calculated as a percentage of all segmented cells; points indicate patient means from tissue replicates, and the solid line depicts linear regression with a shaded 95% CI). Panel **e** maps the mean Z-score of cell type enrichment within a 25 µm radius of cDC2s. Statistical significance was evaluated using a one-sample $t$-test across patients against a null value of 0, with error bars representing the mean ± 95% CI.

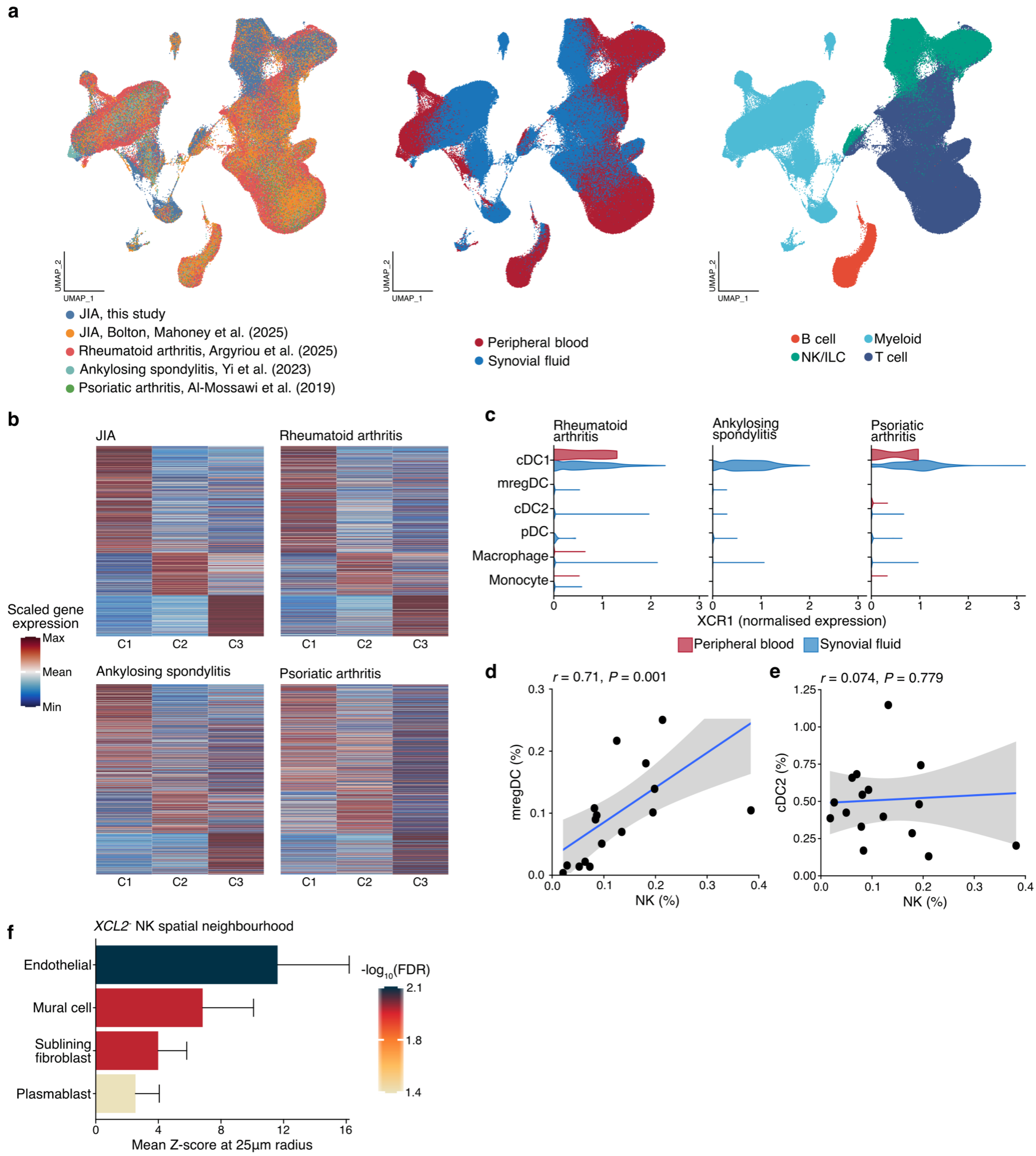


**Extended Data 6: Cross-disease validation of the NK-cDC1 axis in inflammatory arthritis.** **a**, Integrated UMAP comprising public CITE-seq datasets from JIA[1], RA[2], ankylosing spondylitis[3], and psoriatic arthritis[4]. Cells are coloured by study (left), tissue origin (middle), and broad immune cell type (right). **b**, Heatmap displaying row-scaled gene expression profiles that differentiate the NK cell subpopulations C1, C2, and C3, as defined in **Fig. 2c** (CITE-seq). **c**, Normalised *XCR1* expression across annotated myeloid cell populations derived from synovial fluid, separated by disease cohort. **d-f**, Re-analysis of Xenium spatial transcriptomics data of treatment-naïve RA synovial tissue[5] ($n$ = 17 patients). **d**, **e**, Correlation of NK cell abundances with mregDCs (**d**) or cDC2s (**e**), calculated as a percentage of all segmented cells. The solid line represents linear regression, with the shaded region indicating the 95% CI. **f**, Mean Z-scores reflecting cell type enrichment within a 25 μm radius of *XCL2*$^{-}$ NK cells. Statistical significance was determined using a one-sample *t*-test across patients against a null value of 0, with error bars representing the mean ± 95% CI.
